# TIP60, a key lysine acetyltransferase, acts as a wound-induced factor essential for efficient wound response in planaria

**DOI:** 10.1101/2025.09.24.678263

**Authors:** Akansha Pal, Ankit Arora, Bharti Jaiswal, Dasaradhi Palakodeti, Ashish Gupta

## Abstract

Chromatin modifiers are essential regulators of gene expression, DNA repair, replication, and cell division. Among them, histone acetyltransferases (HATs) such as TIP60 play a central role in modulating chromatin dynamics through acetylation of histone and non-histone proteins. TIP60, a member of the MYST family of HATs, is known to regulate key cellular processes, including transcriptional activation, DNA damage response, and cell cycle progression. Although TIP60’s role in stem cell maintenance and differentiation is well established, its function in stem cell-driven regeneration has remained unexplored. In this study, we characterize the role of the TIP60 homolog, SMED-TIP60, in the planarian *Schmidtea mediterranea*, a model organism renowned for its regenerative capacity. Biochemical assays confirmed SMED-TIP60’s acetyltransferase and auto-acetylation activity. RNAi-mediated knockdown of Smed-tip60 resulted in severe defects in tissue homeostasis, survival, and regeneration, including impaired blastema formation and failure to regenerate tissues. In situ hybridization and immunofluorescence analyses revealed a marked reduction in stem cell populations and mitotic activity. Western blotting showed a peak in SMED-TIP60 expression at 5 days post-amputation, suggesting a role during regeneration. RNA-seq analysis revealed widespread dysregulation of gene expression at both anterior and posterior wounds, correlating with increased TIP60 expression post-injury. Notably, wound-response gene expression was aberrant in Smed-tip60 RNAi animals, indicating TIP60’s essential role in initiating wound responses and resetting positional cues. Together, these findings establish TIP60 as a critical regulator of stem cell-mediated regeneration and wound healing in planarians.

## Introduction

TIP60 (Tat-interactive protein 60 kDa), also known as KAT5, is a multi-functional protein, that belongs to the MYST family of lysine acetyl transferases ^1,2^. The name MYST, is an acronym derived from the first letters of the founding members of this family: MOZ, Ybf2/Sas3, Sas2, and TIP60 ^3^. Like other members of the MYST family, TIP60 has distinct functional domains that contribute to its role in gene regulation and chromatin remodelling. These include the chromodomain, which binds to specific histone modifications like methylated lysine residues, a specialized type of HAT (histone acetyltransferase) domain, called as MYST domain found exclusively in the MYST family of proteins, and helps to catalyzes the acetylation of specific lysine residues on both histone and non-histone proteins using acetyl-CoA as the acetyl group donor ^3,4^.

As a histone acetyltransferase, TIP60 can acetylate histones, that facilitates chromatin decompaction making the DNA more accessible for modulating gene transcription ^2,5,6^. The core histones H4 and H3 and H2AX are known substrates of TIP60 ^2,7^. The ability of TIP60 to acetylate non-histone proteins highlights its versatility and critical role in regulating diverse cellular functions beyond chromatin remodelling. TIP60-mediated acetylation of its non-histone substrates can modify the activity, intracellular localization and stability of these proteins, thereby influencing various cellular processes ^8–11^. Due to its critical involvement in key processes such as DNA repair, gene regulation, and maintaining genomic integrity, TIP60 has been extensively studied in humans. It plays a crucial role in safeguarding the integrity of the genome by acetylating and activating ATM kinase and p53 thereby enabling the repair of DNA double-strand breaks (DSBs) and maintaining genomic stability ^4,8,12^. Additionally, TIP60 works as an advisor to cell in the decision-making process between DNA repair and apoptosis, depending on the extent of DNA damage ^12–14^. This prevents the propagation of damaged DNA and ensures that cells maintain their genomic integrity.

Apart from its role in DNA damage repair, apoptosis and gene regulation, TIP60 is vital for the proper functioning and maintenance of stem cells, particularly for the maintenance of hematopoietic stem cells (HSCs), which are responsible for replenishing the blood system throughout an organism’s life ^15^. Similarly, TIP60 plays a crucial role in in maintaining the pluripotency and self-renewal capabilities of mouse embryonic stem cells (ESCs) by binding to active Pol II promoters and a subset of enhancers, thereby regulating gene expression through histone acetylation and interaction with other proteins like c-Myc, which is known to regulate an ESC specific transcriptional module ^15,16^. Recent study by Tominaga K et al emphasized the critical role of TIP60, in maintaining the equilibrium and proper function of neural stem cells (NSCs), required for healthy brain development and function ^17^. Notably, TIP60-p400 complex while promoting self-renewal in ESCs, helps silence the genes that promote differentiation by acetylating histones in a way that keeps certain regions of the genome tightly packed and inaccessible to the transcription machinery ^18^. This way, TIP60 helps maintain a balance by selectively activating and repressing genes and ensures that stem cells can either self-renew or differentiate into specialized cell types as needed. This ability to balance these processes is vital for tissue homeostasis and regeneration.

Other than humans, TIP60 has been studied in several model organisms, including Drosophila, mouse, yeast and zebrafish, providing valuable insights into its functions and mechanisms. In *Drosophila melanogaster* (fruit fly), TIP60 (referred to as dTip60 in Drosophila) has been explored for its roles in chromatin remodelling, DNA repair, and cognition linked processes ^19–21^. Studies in *Mus musculus* (mouse) have extensively investigated TIP60’s functions in DNA repair, apoptosis, and cancer, with knockout mice showing increased sensitivity to DNA damage and a higher propensity for tumor development ^22–26^. Additionally, *Saccharomyces cerevisiae* (yeast) models have been instrumental in understanding the basic mechanisms of histone acetylation and chromatin remodelling by TIP60 ^27–32^. These organisms have contributed significantly to our understanding of TIP60’s diverse and crucial roles in various cellular processes.

Unlike these experimental model organisms, there is limited research on TIP60 in planaria. While planaria are known for their remarkable regenerative abilities, studies focusing on TIP60’s role in these organisms are not widely documented. The simplicity and extraordinary regenerative abilities of planaria have made them an exceptional model organism for studying complex biological processes such as regeneration and stem cell behaviour ^33–35^. The unique regenerative ability of planaria is largely due to their high number of pluripotent stem cells, known as neoblasts, that can turn into any cell type required to rebuild tissues and organs ^36–38^. When a planaria is cut into pieces, the neoblasts in each fragment migrate to the wound site and start dividing and differentiating into the necessary cell types to form a complete, new fully functioning worm. The self-renewing ability of neoblasts allows these cells to divide and produce more stem cells, which is essential for continuously replenishing the stem cell pool for tissue maintenance and regeneration ^38,39^. The presence of neoblasts throughout the planaria’s body ensures that any injury can be quickly and efficiently repaired, regardless of where it occurs.

Regeneration is indeed a multifaceted and highly complicated process that involves a series of intricate and well-coordinated biological processes, including the proliferation of neoblasts, the migration and differentiation of neoblasts to form new tissue, programmed cell death to remove old or damaged cells, and autophagy for recycling cellular components ^36,39,40^. Many signalling pathways like Wnt/β-catenin, Hedgehog, and Notch, along with apoptosis regulators, autophagy-related proteins, transcription factors, and epigenetic modifiers, coordinate tissue patterning and gene regulation during regeneration ^41–49^. Despite significant progress, our understanding remains incomplete, and many uncharacterized factors await discovery to fully elucidate the intricacies of regeneration. In this study, we characterized the planarian homolog of the human TIP60 protein. To study and investigate the process of TIP60-mediated gene regulation in planaria and further advance our understanding of the fundamental processes like tissue repair and regeneration in a simpler organism like planaria, it is imperative to biochemically and functionally characterize the homolog of this protein, in detail. This characterization would allow us to extrapolate its properties from the well-studied human counterpart, thus providing deeper insights into its functional roles. Unlike other model organisms such as salamanders, which face limitations in fully elucidating the mechanisms of whole-body regeneration, planaria present a promising model. Because they are capable of whole-body regeneration, they facilitate *in vivo* studies and are crucial for advancing our understanding of the function of TIP60 *in vivo*.

To gain insights into the fundamental functions of these proteins, in planaria, we conducted a comprehensive analysis which included comparisons of sequence homology, domain architecture, phylogenetic analysis, structural similarities and differences, and alongside an examination of biochemical properties, with the human counterpart. From our findings, we can infer that the planarian homolog of TIP60 shares crucial structural and functional features with its human counterpart, despite having limited overall sequence similarity. The catalytic domain demonstrated significant resemblance both in sequence and structure to the human TIP60. The conservation of key functional domains ensures the preservation of essential biochemical activities across different species. The planarian TIP60 exhibited *in vitro* acetylation and autoacetylation activity just like their human counterparts. The knockdown of *tip60* significantly affects their ability to regenerate and maintain homeostasis. To understand the regulatory role of *Smed-tip60* during regeneration, we performed RNA-Seq analysis of *tip60* RNAi animals at 2 DPA and 5 DPA (Days post-amputation). TIP60 depletion led to distinct transcriptional changes at anterior and posterior side, suggesting region-specific roles. Additionally, altered expression of several wound-induced genes, including ectopic and prolonged *wnt1* expression, highlights TIP60’s role in modulating early wound responses and polarity during regeneration.

## Material and Methods

### Animal Maintenance

The sexual strain of the planarian *Schmidtea mediterranea* (S2F2) was utilized in this study. These organisms were maintained in a 1X Montjuïc salt solution at a constant temperature of 20°C. To sustain the stock culture, the planarians were fed goat liver paste twice weekly. Before any experimental procedures, the planarians were subjected to a fasting period of at least one week.

### *In silico* structure modelling and phylogenetic analysis

The sequence of the human TIP60 protein (Q92993) and *S. cerevisiae* (Q08649) **(Supp. figure 1 A)** was obtained from UniProt. Human TIP60 protein sequence was then used to perform a tBLASTn against the protein database of *S. mediterranea* as available on the PlanMine database ^50^. BLASTp hits with a bit score of less than 100 and hits with incomplete protein sequences were removed before further analysis. The homolog of SMED-TIP60 was annotated based on percentage similarity with human TIP60 protein and bit score. Sequence similarity and identity were analyzed by pairwise sequence alignment using the EMBOSS Needle tool. We also did the reciprocal BLASTp as described in to eliminate any false positive hit ^51^. Domains were annotated by using the hmmscan tool (HMMER) ^52^ using the Pfam database ^53^. Domain maps were generated using the IBS Illustrator for biological sequences ^54^.

Three-dimensional models of human, *S. mediterranea*, and *S. cerevisiae* TIP60 protein were generated using homology modelling. To achieve this hsTIP60, ScESA1, and SMED-TIP60 proteins full-length sequences, the chromodomain and MYST domain sequences were used as templates to generate structures of the respective queries using the comparative modeling tool on the Robetta server ^55^. All parameters were set to default. The server predicted 5 best models for each query sequence. These predicted models were checked for stereochemical properties using the SAVESv6.0 server. The model that showed the highest quality for these parameters was further refined using the GalaxyRefine tool ^56^. RMSD analysis of proteins was done using PyMOL ^57^.

For phylogenetic analysis, the human TIP60 protein sequence was used to perform BLASTp (NCBI) against the Refseq protein database model organisms from different phyla for instance Animals, Plants, Fungi, red algae, and green algae. The obtained full-length protein sequences were aligned using the Clustal Omega algorithm ^58^ available on MEGA-X tool ^59^. The alignment was used to predict the best substitution model for phylogeny tree construction^60^.

### RNA extraction, cDNA synthesis and cloning

Total RNA was extracted from the planarians using RNAiso Plus reagent (Takara cat. No. 9108). Approximately 10 medium-sized planarians were incubated in 500µl of RNAiso Plus reagent and stored at −80°C for at least one hour. The animals were then thawed on ice and homogenized thoroughly using a micro pestle (or vortex). Following homogenization, 1/5th volume of ice-cold chloroform was added and mixed vigorously. The mixture was kept on ice for 5-10 minutes and then centrifuged at 12,000 rpm for 30 minutes at 4°C. The top aqueous phase was collected, transferred to a new micro centrifuge tube, and mixed with an equal volume of ice-cold 100% isopropanol. After proper mixing, the mixture was incubated at −20°C for 30 minutes to 1 hr, followed by centrifugation at 12,000 rpm for 30 minutes at 4°C. A black RNA pellet was obtained, after which the supernatant was carefully removed. The pellet was washed twice with 75% ethanol, air-dried, and subsequently resuspended in nuclease-free water.

The extracted RNA was used for cDNA synthesis and can also be stored long-term at −80°C. 1µg of RNA was used to synthesize cDNA using a SuperScript III Reverse Transcriptase cDNA synthesis kit (Invitrogen 18080093), following the manufacturer’s protocol. The *Smed-tip60* gene sequence was obtained from PlanMine (http://planmine.mpibpc.mpg.de/) and specific primers were synthesized. The *Smed-tip60*-specific primers used in this study are mentioned in the **(Supp. table 1).** The *Smed-tip60* gene was amplified from cDNA using Taq polymerase and amplified *Smed-tip60* amplicon was then cloned into the pET28(a) vector (containing a His-tag) using BamHI and SalI restriction sites for recombinant protein purification. Further

### Recombinant protein purification

To purify the recombinant protein, BL21(DE3) Codon Plus cells were transformed with the pET28(a)-*Smed-tip60* plasmid and cultured at 37°C.Once the culture reached an optical density of 0.6, protein expression was induced by adding Isopropyl-β-D-thiogalactopyranoside (IPTG) to a final concentration of 1 mM, followed by incubation at 20°C for 15 hours. The induced culture was then pelleted, and the cell pellet was lysed using an ice-cold lysis buffer. After induction, the cells were harvested by centrifugation, and the resulting pellet was lysed using an ice-cold lysis buffer. Pre-equilibrated Ni-nitrilotriacetic acid (Ni-NTA) beads were added to the lysate, and the mixture was incubated with constant rotation at 4°C for 1 hour. The beads bound to the target protein were collected by centrifugation and washed three times with lysis buffer containing 20 mM imidazole. The His-tagged proteins were then eluted using lysis buffer with 500 mM imidazole for 30 minutes at 4°C. Finally, the eluted proteins were dialyzed in a buffer containing 50 mM Tris-Cl (pH 8.0), 2 mM EDTA, 100 mM NaCl, and 0.2 mM PMSF, and the dialyzed protein was stored in aliquots at −80°C.

### Histone Acetyltransferase (HAT) assay and Autoacetylation assay

HAT activity was assessed by using recombinant His-SMED-TIP60 protein. The HAT reaction was performed in a total volume of 50 μL, containing dialyzed His-SMED-TIP60 protein, 0.5 μg of histone H4 peptide (Millipore), 100 μM acetyl coenzyme A (Sigma), and HAT assay buffer (50 mM Tris-Cl [pH 8.0], 10% glycerol, 0.1 mM EDTA, 1 mM DTT and 0.2 mM PMSF). Reactions were incubated at 30 °C for 1 hour and terminated by adding 2× Laemmli sample buffer. Samples were then boiled at 95 °C for 10 minutes, resolved by 15% SDS-PAGE and analyzed via Western blotting using anti-acetylated histone H4 antibody (1:1,000; Millipore) and to histone H4 antibody (1:1,000; Millipore).

To assess autoacetylation, purified His-SMED-TIP60 protein was incubated with 100 μM acetyl coenzyme A in HAT buffer for 1 hour at 30 °C. The reaction was stopped by the addition of 2× Laemmli sample buffer. Samples were then boiled at 95 °C for 10 minutes and subjected to 10% SDS-PAGE. Western blotting was performed using an anti-acetylated lysine antibody (1:1,000; Cell Signaling Technology).

### Yeast complementation assay

The *Saccharomyces cerevisiae* temperature-sensitive ESA1 mutant strain LPY3500 was obtained as a generous gift from Dr. Lorraine Pillus (University of California at San Diego, La Jolla, CA). The strain was transformed with plasmid constructs pRS314 (empty vector), pRS314-Sc-esa1 (encoding the yeast *ESA1* gene), and pRS314-*Smed-tip60* (encoding *Smed-tip60*) using the lithium chloride transformation method. Transformed cells were selected on dropout media lacking tryptophan and incubated at 25 °C for 3–4 days to allow colony formation. To assess the complementation efficiency of the introduced constructs, a master plate was prepared by streaking individual transformants onto tryptophan-dropout media plates. Colonies were subsequently replica-plated onto fresh-Trp plates and incubated under both permissive (25 °C) and restrictive (37 °C) temperature conditions. Growth at 37 °C was used as an indicator of functional complementation by the expressed construct.

### dsRNA and RNA Probe Synthesis

Gene-specific primers with a range of 600-700bp were used to amplify the *Smed-tip60* **(Supp. table 1)**. Taq Polymerase enzyme (TaKaRa RR002C) was used for the PCR amplification. The PCR amplicon was purified using the PCR cleanup kit (Promega kit A9281). Purified PCR product was cloned in the Topo vector using the TA cloning kit followed as mentioned in the manufacturer’s protocol (Invitrogen K207040). The ligated product was used to transform in DH5α cells and screening for blue-white colonies was done for generated clones under the kanamycin selection media. The gene-specific primers, with the T7 promoter overhang were used to amplify the gene using the Taq polymerase enzyme. An *in vitro* transcription reaction was set up using the T7 promoter overhang templates to produce dsRNA. For probe synthesis, a DIG RNA labelling mix was used for *in vitro* transcription reaction.

### RNAi microinjection

*Smed-tip60* RNAi was performed using the microinjection method as described in ^61^. Purified dsRNA of *Smed-tip60* and Green Fluorescent Protein (GFP) (used as a negative control) with a final concentration of 2µg/µl was used. Animals were immobilized under a stereo microscope (Olympus SZ61) on a petri dish filled with ice and injected with 90 nl of dsRNA using a Nanoject II Drummond microinjector. Injections were administered every alternate day for seven times. For regeneration studies, animals were amputated into three pieces (head, trunk, and tail) 24 hours after the last microinjection. These animals were fixed and used for further studies.

### Fixation and fluorescence *in situ* Hybridization (FISH)

FISH was performed following protocol as described elsewhere ^62^. Animals were incubated in 5% N-Acetyl Cysteine (prepared in 1X PBS) for at least 5 minutes and fixed in 4% formaldehyde (prepared in PBSTx) for 15 minutes. The animals were then washed twice with PBSTx, dehydrated in a methanol gradient, and stored at −20°C in methanol. For bleaching, fixed animals were rehydrated and incubated in a bleaching solution of 5% formaldehyde, 1.2% H_2_O_2_, and 0.5x SSC under a light source for approximately 3 hours. After bleaching animals were washed with PBSTx, treated with 2µg/ml proteinase K, and fixed again in 4% formaldehyde in PBSTx. Animals were washed in 1:1 PBSTx: Prehybridization solution followed by incubation in prehybridization buffer for 2 h at 56°C. Next, RNA probes were mixed in reducing hybridization buffer with a final concentration of 0.5ng/µl and added to the samples, and incubated for at least 18h at 56°C. Subsequently, the samples were washed using TNTx buffer and incubated in a blocking solution (5% horse serum and 5% Roche western blocking reagent) for 2h at room temperature. This was followed by an overnight incubation in anti-Dig POD (1:1000) at 4°C and then signal was developed using the tyramide signal amplification method.

### Immunostaining

Animals were fixed in 4% formaldehyde and bleached as described above. After bleaching, the animals were incubated in a blocking solution containing 10% horse serum in PBSTx. Primary anti-H3P antibody (Invitrogen cat. no. PA5-17869, 1:100) prepared in blocking solution was added to the animals and incubated for 48 hours at 4°C. For nuclei staining, DAPI (1:1000 from 5 mg/ml stock; Sigma-Aldrich [cat. no. D9542]) was used.

### RNA sequencing and transcriptomic analysis

Bulk RNA was extracted at 2 DPA and 5 DPA from the anterior and posterior side of trunk blastema for control and *Smed-tip60* knock down animals. The experiment was conducted in three biological replicates. Poly A selection was performed using the NEBNext Poly(A) mRNA Magnetic Isolation Module (Catalog no. E7490L), and transcriptome library preparation was carried out with the NEBNext® Ultra™ II Directional RNA Library Prep Kit with Sample Purification Beads (Catalog no. E7765L). Sequencing was conducted on the NovaSeq 6000 platform utilizing an SP flow cell with a paired-end read length of 2×50 bp. Raw RNA-seq reads underwent quality control and trimming using fastp [v0.20.1] ^63^, and the resulting QC reports were compiled using MultiQC [v1.9] ^64^. Read alignment was performed with STAR [v2.7.9a] ^65^ against the S. mediterranea (S2F2) reference genome. Aligned reads were then used to generate raw gene expression counts with featureCounts ^66^ from the Subread package [v1.5.2]. Differential gene expression was analyzed for differential expression, using DESeq2 [v1.30.1] ^67^ in R environment [v4.0.3].

### Single cell RNA sequencing data analysis

UMI counts in form of expression matrix were obtained from Fincher et. al 2018, Zeng et. al 2018, Raz et. al 2021 and Scimone et. al 2022 were analysed using Seurat (v4.0.0) ^68^ in R environment (v4.0.3). Normalization on cells was performed using LogNormalize normalization method and cells were filtered for genes expressing at least 200 cells. After normalization, genes were used as input for executing principal component analysis (PCA) using function RunPCA and genes were used for visualization 2D data in Uniform Manifold Approximation and Projection (UMAP) form using function RunUMAP. In Seurat, clustering was performed using the ‘FindClusters’ function adopting the resolution of 0.1 and the shared-nearest-neighbor (SNN) graph were constructed using the first 30 PCA dimensions.

### Microscopy and image analysis

Brightfield and darkfield imaging were performed using an Olympus SZX16 microscope. Confocal images were acquired using an FV3000 laser-scanning microscope and processed and quantified using Fiji (version-ImageJ2.1. o/1.53c). Graph Pad Prism (version-8.0.1) was used for graph plotting and statistical analysis.

## RESULTS

### *S. mediterranea* TIP60 predominantly expresses in neoblast cells

For comparative analysis of TIP60 homologs, we searched the *S. mediterranea* genome ^69^ for SMED-TIP60 **(dd_Smed_v6_4706_1_2)** and included the *S. cerevisiae* TIP60 homolog (ScEsa1) and HsTIP60 as a reference in our analysis. Multiple sequence alignment conducted with the protein sequences of HsTIP60, ScEsa1, and SMED-TIP60, combined with domain annotation from hmmscan tool (HMMER 3.4)^52^ using Pfam database ^53^, revealed specific conserved domains in SMED-TIP60 suggesting that these proteins may have similar functions (**Supp. figure 1A**). The domain mapping, illustrated in **Figure 1A**, indicated that SMED-TIP60 has a conserved chromodomain ranging from **5-60** amino acids, a zinc finger domain spanning **170-224** amino acids, and a MYST domain from **229-444** amino acids. The percent identity matrix indicated high conservation of the catalytic MYST domain, with **61.6%** identity to the human homolog and **54.2%** identity to the yeast homolog.

**Fig. 1.**
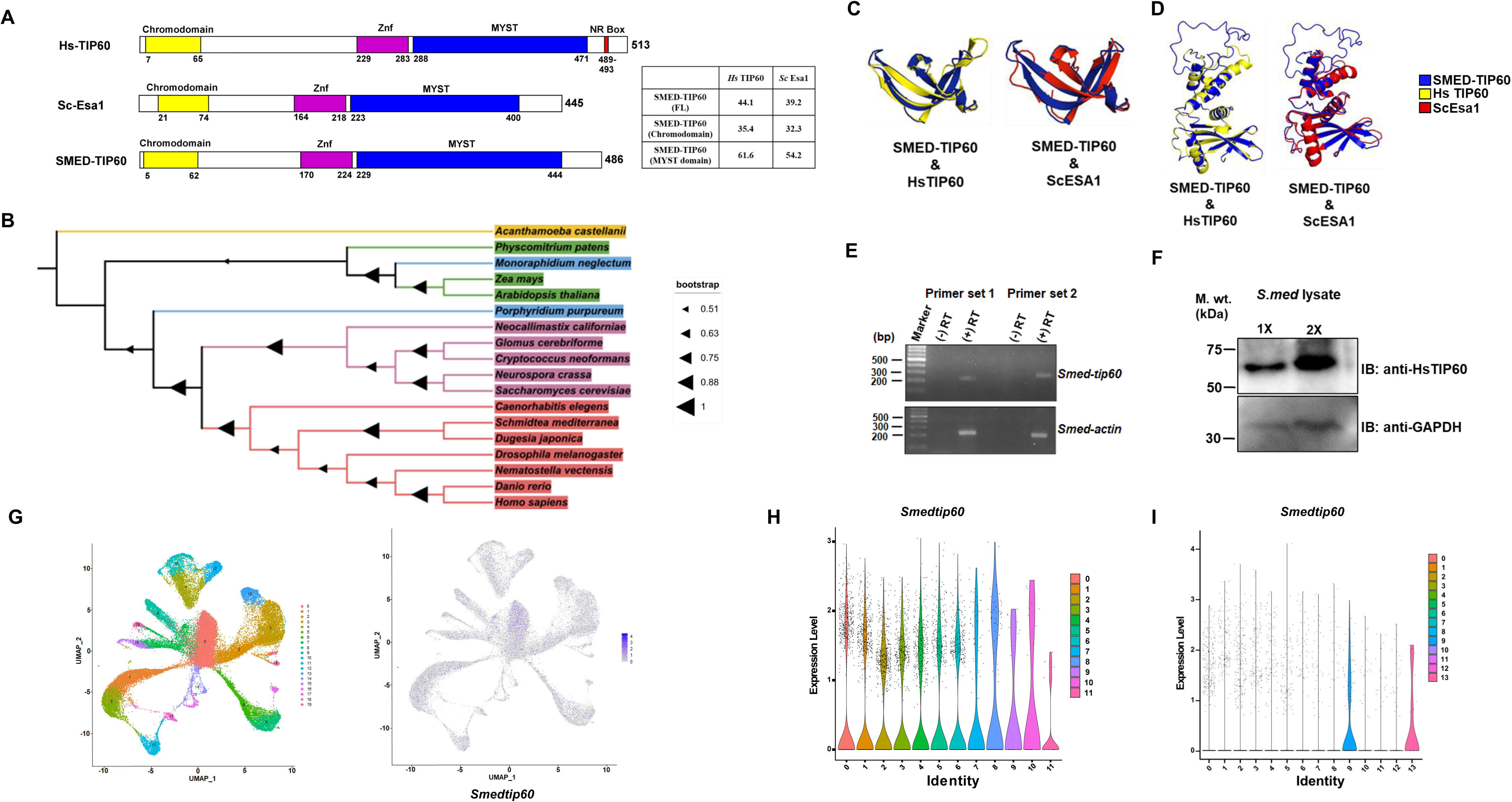
Characterization of SMED-TIP60 and its expression in planaria. (A) Comparative domain analysis of TIP60 protein of *humans*, *Saccharomyces cerevisiae* (ScESA1), and *Schmidtea mediterranea* using HMMER and Pfam databases, highlighting conserved domains. The table shows the percentage identity between full-length proteins and conserved MYST domain. (B) The phylogenetic tree for the TIP60 proteins of model organisms from different kingdoms was constructed using MEGA X and visualized on iTOL software. Color coding is according to Kingdom like Protista in yellow and blue color, Fungi in Purple color, Plantae in green color, and Animals in Red color (C) Superimposition of SMED-TIP60 (Blue color) and HsTIP60 (Yellow color) Chromodomain 3-D modelled structures shows RMSD of 0.514Å and SMED-TIP60 (Blue color) and ScESA1 (Red color) shows RMSD of 2.005Å. (D) Superimposition of 3-D modelled structure of MYST domain for SMED-TIP60 (Blue color) and HsTIP60 (Yellow color) shows RMSD of 0.049 Å, SMED-TIP60 (Blue color) and ScESA1 (Red color) shows RMSD of 0.447 Å. (E) PCR products obtained using two primer sets (Set1 and Set2) were resolved on agarose gel. (+) RT and (-) RT lanes indicate the PCR product obtained using a control template generated with and without reverse transcriptase enzyme. *Smed-actin* was amplified using two primer sets (set1 and set2) from the same samples. (F) Endogenous SMED-TIP60 protein was detected by Western blot. *S.med* lysate in two different concentrations was resolved on SDS-PAGE followed by Western blot analysis using the HsTIP60 antibody and anti-GAPDH was performed as a loading control. (G) UMAP plot showing total number of clusters obtained in scRNA-seq dataset of Whole animal in planaria from Fincher et al., 2018. Similarly, feature plot shows the *Smed-tip60* expression in obtained clusters. (H) Violin plot exhibiting the *Smed-tip60* expression in scRNA-seq dataset across X1 cell population clusters (Zeng et.al., 2018) (I) Violin plot showing *Smed-tip60* expression in scRNA-seq profiles from the cells in the G0/G1 phase of X2 population (Raz et al., 2021).

We expanded the study to include a phylogenetic analysis to understand the evolutionary relationships and diversity of the TIP60 protein among various taxonomic kingdoms, including representatives from Protists (*Acanthamoeba castellanii*, *Monoraphidium neglectum*, *Porphyridium purpureum*), Fungi (*Neocallimastix californiae, Glomus cerebriforme, Cryptococcus neoformans, Neurospora crassa, Saccharomyces cerevisiae*), Plantae (*Physcomitrium patens, Zea mays, Arabidopsis thaliana*), and Animalia (*Caenorhabditis elegans, Schmidtea mediterranea, Dugesia japonica, Drosophila melanogaster, Nematostella vectensis, Danio rerio, Homo sapiens*) **(Supp. table 2)**. Using the blastp method, we identified homologs of the human TIP60 protein (UniProt accession Q92993) from a broad array of model organisms representing different kingdoms. Multiple sequence alignments were conducted for TIP60 protein sequences from these organisms, followed by constructing a phylogenetic tree using the maximum likelihood method in MEGA (Molecular Evolutionary Genetics Analysis) software ^59^. The analysis revealed clear separation of the examined organisms into three major clades: Plantae, Fungi, and Animalia, supported by high bootstrap values, indicating significant evolutionary divergence of the TIP60 protein among these taxa aligned with the respective organismal complexity and ecological niches **(Figure 1B)**. The homolog identified in planaria clusters with homologs from *humans, C. elegans, Danio rerio, Drosophila, Dugesia, and Nematostella.* This indicates that the protein is conserved across these diverse species, suggesting it has an essential function that has been preserved throughout evolutionary history in the animal kingdom.

To further elucidate functional and evolutionary conservation, we constructed 3D model using Robetta webserver ^55^, and conducted a full-length structural comparison of SMED-TIP60 to HsTIP60 and ScEsa1 proteins **(Supp. figure 1B)**. RMSD (Root Mean Square Deviation) analysis was employed to assess structural variations. The RMSD value between SMED-TIP60 and HsTIP60 was 16.298 Å, indicating substantial dissimilarity in their full-length structures **(Supp. figure 1B, bottom panel)**. The RMSD value between SMED-TIP60 and ScEsa1 was 10.505 Å, further highlighting distinct structural differences among these proteins. In addition to modelling the full-length protein structures, we also conducted an analysis of the chromodomain and MYST domain, which corresponds to the catalytic domain. This analysis revealed significantly low RMSD values for the catalytic domains of SMED-TIP60 and HsTIP60 indicating a high degree of structural conservation in this region **(Figure 1C and 1D)**. These findings suggest that despite the structural differences in the full-length protein structures of SMED-TIP60 with human, the preservation of the catalytic domain implies the potential conservation of the protein’s functional role. The combined data from phylogenetic and structural analyses suggest that TIP60 homologs have undergone significant evolutionary divergence, resulting in both conserved and specialized features across different kingdoms.

To examine the endogenous expression of SMED-TIP60 in planaria, we conducted a semi-quantitative PCR analysis using the SMED-TIP60 sequence retrieved from the PlanMine server ^50^ (**dd_Smed_v6_4706_1_2**). Two sets of primers were designed: primer set 1 from the mid-region and primer set 2 for the C-terminal of the gene, as depicted in **Supp. figure 1C**. Our PCR results revealed the presence of *Smed-tip60* transcripts, demonstrated by the specific band size observed in the agarose gel electrophoresis image **(Figure 1E)**, and confirms that the *tip60* gene is transcribed in *S. mediterranea*. Furthermore, to verify that these transcripts are translated into functional protein, we performed a protein expression analysis using a human TIP60 antibody, which cross-reacts with the SMED-TIP60 protein **(Figure 1F)**. The detection of the TIP60 protein validates that the gene is both transcribed and translated in the planaria system. To investigate the cell or tissue specific expression profile of SMED-TIP60 in planaria, we employed publicly available single-cell RNA sequencing (scRNA-seq) datasets for high-resolution analysis and identification of SMED-TIP60 in specific cell populations expressing the planarian homolog. We re-analyzed scRNA-seq dataset obtained from fincher et. al. 2018 in whole animal to extract and check for the cells which express *Smed-tip60* ^70^. We found 6351 cells (out of 50,562 cells as shown in UMAP plot) while feature-plot expressing *Smed-tip60* transcripts in the single-cell transcriptome (**Figure 1G**). In general, the expression S*med-tip60* is lower across the clusters and among different clusters, neoblast cluster (Cluster 0) showed the maximum expression. Reconstruction of the violin plot from the ScRNA-seq obtained from X1 population ^71^ showed expression of *Smed-tip60* in cluster 0, 1, 2, 3, 4, 5, 6, 7, 8, which corresponds to clonogenic, epidermal, gut, muscle and pharyngeal neoblast population suggesting that SMED-TIP60 might play an essential role in maintenance of neoblast and its progeny **(Figure 1H).** Furthermore, transcriptome analysis of scRNA-seq datasets from Raz et. al 2021 from X2 population (G0/G1) showed weak expression of *Smed-tip60* in cluster 9 and 13 ^72^ **(Figure 1I).** We also analysed the scRNA-seq data from Scimone et. al 2022 generated from the anterior and posterior cut regions at 0 HPA, 18 HPA ^73^. Our analysis revealed high expression of *Smed-tip60* at 0 HPA, 18 HPA-anterior and 16 HPA-posterior suggesting the potential role of SMED-TIP60 in wound healing during regeneration **(Supp. figure 1D).**

### Smed-TIP60 possess histone acetyl transferase activity and autoacetylation activity

Given that lysine acetyltransferases (KATs) are enzymes responsible for transferring acetyl groups from acetyl-CoA to lysine residues of substrate proteins ^74–76^, we aimed to biochemically characterize the SMED-TIP60 protein to determine if it functions as a lysine acetyltransferase, like its human counterpart. For this we conducted an *in vitro* HAT assay using the recombinant SMED-TIP60 protein. For purification of recombinant SMED-TIP60 protein, we first cloned the SMED-TIP60 ORF encoding the SMED-TIP60 protein into the pET-28a expression vector and validated positive clones through restriction enzyme digestion **(Supp. figure 2A)**. For expression and purification of recombinant SMED-TIP60 protein, BL21(DE3) Codon Plus bacterial cells were transformed with the pET28a-SMED-TIP60 clone, followed by protein induction with IPTG. Western blot analysis confirmed successful induction of the His-tagged recombinant SMED-TIP60 protein (**Figure 2A**). The expressed protein is then purified using Ni-NTA affinity chromatography under native conditions **(Figure 2B)**. Recombinant SMED-TIP60 protein was then subjected to *in vitro* HAT assay as described in the method section. The transfer of acetyl groups to lysine residues on the substrates was determined through Western blotting using specific acetyl-lysine antibodies **(Figure 2C)**, which clearly show that recombinant SMED-TIP60 can acetylate histone H4 peptides *in vitro*. To investigate whether the SMED-TIP60 protein possesses autoacetylation sites, we did pairwise sequence alignment with the human TIP60 protein sequence and marked the putative sites of autoacetylation in SMED-TIP60 protein **(Supp. figure 2B)**. Further, we performed *in vitro* autoacetylation assay, a reaction set up with SMED-TIP60 protein and acetyl CoA at 30° C. Western blot analysis with lysine antibodies confirmed that SMED-TIP60 exhibits autoacetylation activity **(Figure 2D)**.

**Fig. 2.**
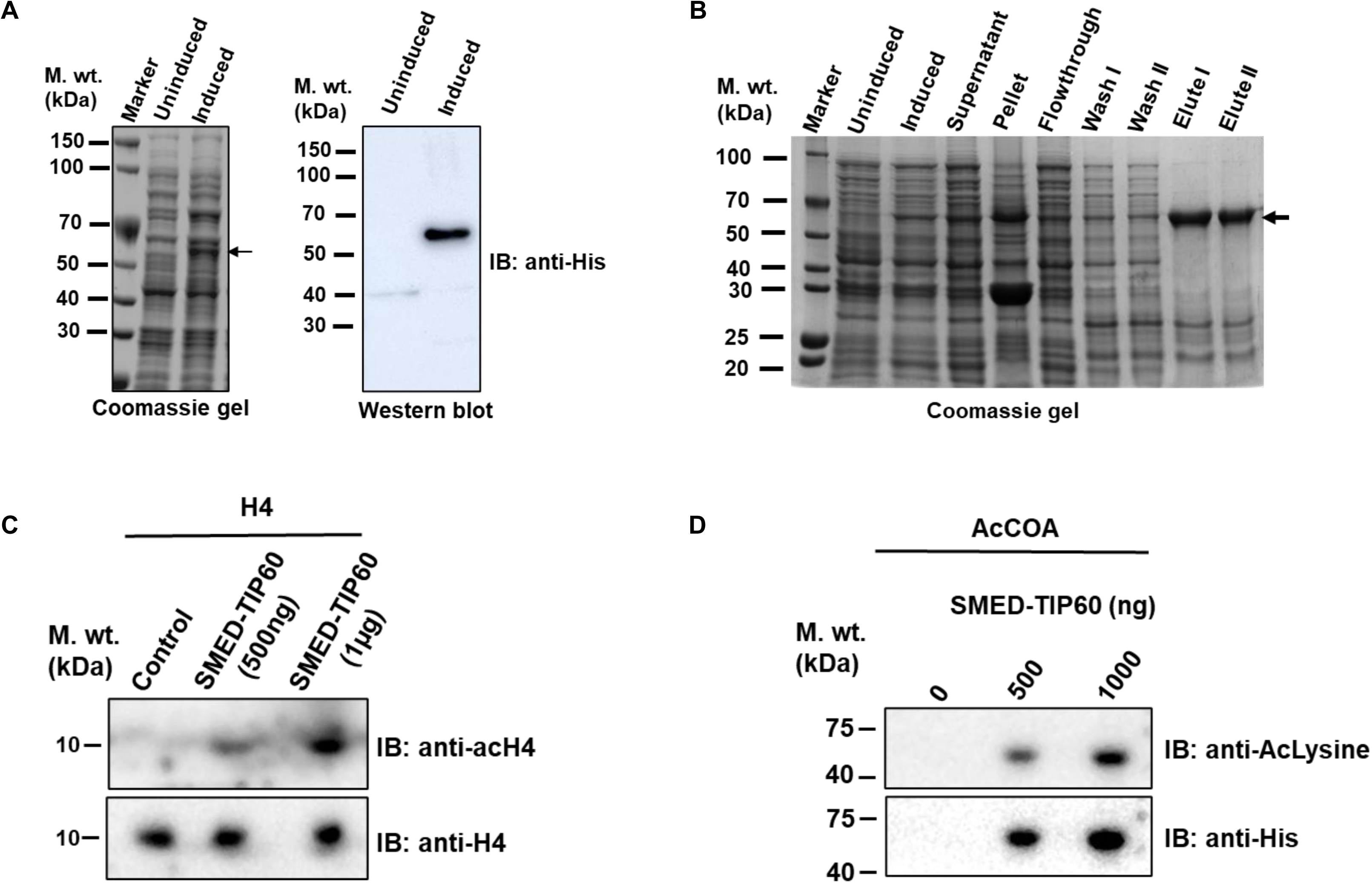
Smed-TIP60 possesses acetyl-transferase activity. (A) Expression of recombinant SMED-TIP60 protein. BL21 DE3 (codon plus) cells were transformed with pET (28)a-*Smed-tip60* plasmid and used IPTG to induce the recombinant SMED-TIP60 protein expression. Lysates from uninduced and induced samples were resolved in Coomassie gel. Arrow showed expression of induced His-SMED-TIP60 protein. Western blot was performed from the same samples using an anti-His antibody, showing a signal at the expected size of ∼55KD in the induced lane. (B) The purified protein expression profile of His-SMED-TIP60 was resolved in Coomassie gel. A band of His-SMED-TIP60 protein was observed at ∼55kDa marked with the arrow. (C) SMED-TIP60 can acetylate H4. *In vitro* Histone acetyltransferase assay used recombinant His-SMED-TIP60 protein and Histone 4 peptide as substrate. The reaction mix was resolved on 15% SDS-PAGE followed by Western blot analysis with indicated antibodies. (D) *In vitro* autoacetylation assay was performed using different concentrations of recombinant SMED-TIP60 protein. The samples were resolved on 10% SDS-PAGE followed by Western blot. The blots were probed with an anti-aclysine antibody and an anti-His antibody.

To determine whether the functional role of the *S. mediterranea* TIP60 protein is conserved across species, we performed a yeast complementation assay, which are typically done to determine whether a gene from one organism (in this case, *Smed-tip60*) can compensate for the loss or malfunction of a corresponding gene in another organism (in this case, *S. cerevisiae* ScEsa1). For this purpose, we cloned the *S. mediterranea tip60* gene into the pRS314 yeast expression vector **(Supp. figure 2C)** and subsequently transformed the pRS314-*Smed-tip60* construct or pRS314-ScEsa1 or pRS314 vector, into *S. cerevisiae* LPY3500 strain, which harbours a mutation in the HAT domain of the *ScEsa1* gene, resulting in cell cycle progression defects at restrictive temperatures ^77^. The wild-type ScEsa1 gene was able to rescue the cell growth defect under these conditions. However, the *Smed-tip60* gene failed to complement this function, indicating that the TIP60 protein from *S. mediterranea* does not substitute for the function of the ScEsa1 protein in the yeast system **(Supp. figure 2D)**. This suggests that there may be species-specific differences in the functional roles or interactions of TIP60 protein. Together these findings including SMED-TIP60’s ability to acetylate histone and exhibit autoacetylation activity, suggest its functional similarity to its human counterpart. Despite these *in vitro* similarities, SMED-TIP60 does not complement the function of ScEsa1 in the yeast system, indicating functional divergence.

### Knockdown of SMED-TIP60 induces lethality and regeneration defects in planaria

To decipher the role and significance of SMED-TIP60 protein in planarian homeostasis we performed RNA interference (RNAi) analysis and assessed the impact of *tip60* gene loss by thoroughly observing the phenotypes of intact planarians, to discern any potential alterations or abnormalities in behaviour and morphology resulting from the knockdown. For RNAi experiments, dsRNA of ∼600bp complementary to the mRNA of the *tip60* gene, was synthesized and introduced by directly injecting into the planaria, using a microinjector ^61^. To ensure consistence presence of the *tip60*-dsRNA in the worm, and achieve sustained knockdown of the *tip60* gene, a regime of seven injections at one-day intervals was followed **(Figure 3A)**. To verify the knockdown efficiency, qPCR and Western blot was performed, using lysates from control and *Smed-tip60* knockdown (KD) animals. The result showed a significant reduction in both mRNA and protein levels compared to controls, indicating effective knockdown **(Figure 3B, Supp. figure 3A)**. Following the injections, we assessed the animals for any phenotypic defects in comparison to control. Interestingly, we observed that *Smed-tip60* KD animals displayed initial signs of head regression, which gradually progressed to head lysis **(Figure 3C)**, eventually leading to the complete lysis of all animals within a 100-day period **(Figure 3D)**. This severe sequence of events indicates the crucial role of TIP60 in maintaining cellular integrity and tissue organization.

**Fig. 3.**
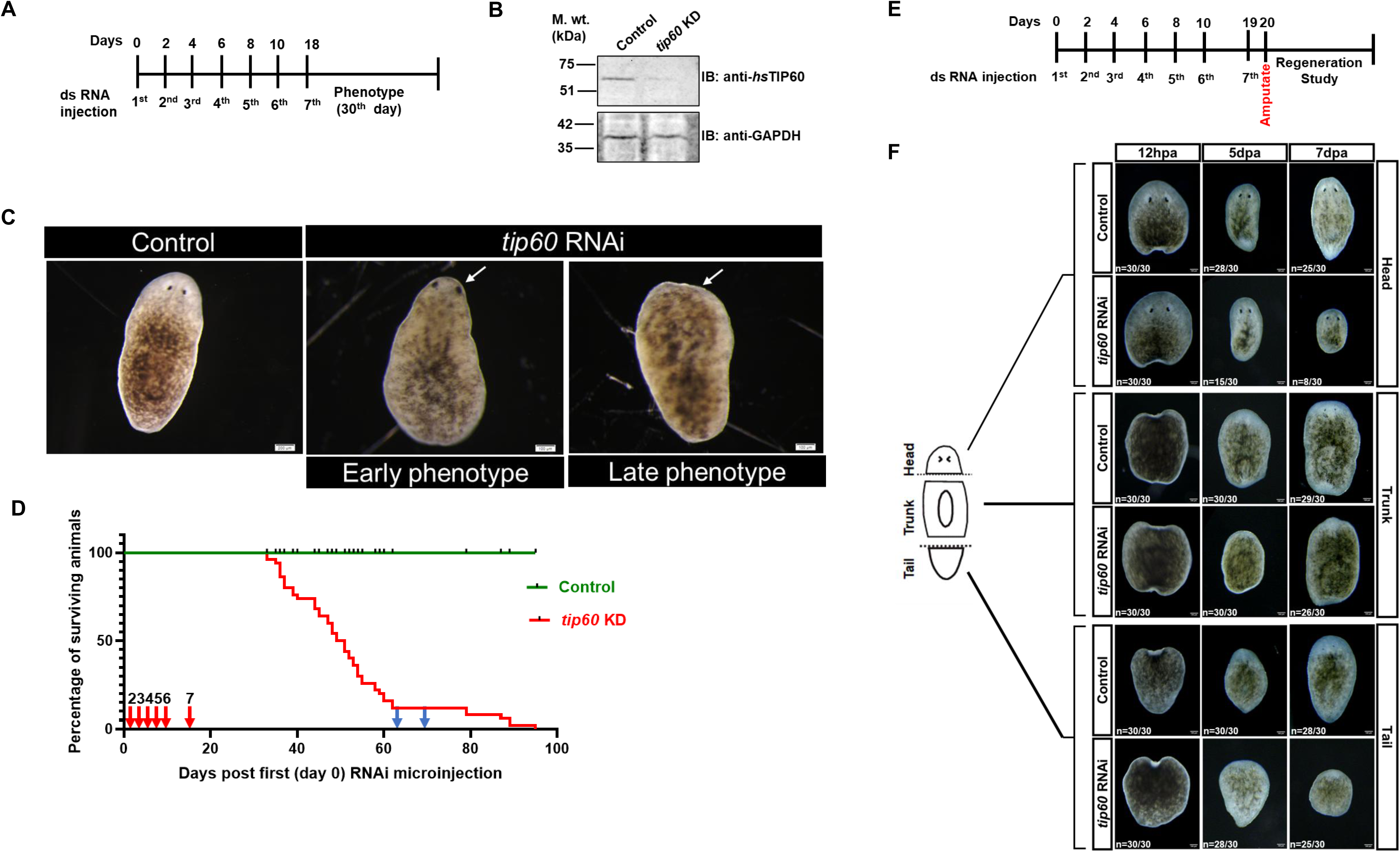
SMED-TIP60 is required to maintain homeostasis and regeneration in planaria. (A) The regime followed for control and *Smed-tip60* dsRNA microinjection procedure during homeostasis condition. (B) Lysates were prepared from control and *Smed-tip60* RNAi planarians. Samples were resolved on SDS-PAGE gel followed by Western blot analysis using antibodies indicated above. (C) Dark-field images of intact planarians under control and *Smed-tip60* RNAi condition. Images showing early and late phenotypes, and white arrows indicate head regression and head lysis in the *Smed-tip60* RNAi condition. (n=30, scalebar = 200µm). (D) Survival plot illustrating the percentage of intact planarian’s survival after microinjection with control and *tip60* dsRNA, with red arrows indicating dsRNA microinjection. The first microinjection was given from day 0 and blue arrows indicated subsequent dsRNA feedings n= 50. (E) The regime followed for dsRNA microinjection during regeneration studies planarians were amputated transversely after 24 hours of the last microinjection. Bottom panel: Dark-field images of control and *Smed-tip60* RNAi planarians during the regeneration time course from 12 HPA, 5 DPA, to 7 DPA. (Note: HPA refers to hours post-amputation and DPA - Days post-amputation).

Considering the pronounced effects observed on the heads of intact planarians following gene knockdown, characterized by regression and eventual mortality, we wanted to investigate the impact of TIP60’s absence on their regenerative capabilities. Following a similar injection regime as previously described, the planarians were transversely amputated into three segments-head, trunk, and tail-24 hours after their final injection (**Figure 3E**). Notably, we observed that RNAi-mediated disruption of *tip60* led to significant defect in the regeneration process, particularly the inability to form a blastema, a critical initial step in the regenerative process. Even at 5 days post-amputation (DPA), the knockdown animals failed to form a proper blastema (**Figure 3F**). This inability to form a blastema upon *tip60* RNAi knockdown indicates that TIP60 may play an essential role in the early cellular and molecular events required for regeneration. Together, these findings demonstrate that SMED-TIP60 is essential for both maintaining homeostasis and facilitating regeneration.

### SMED-TIP60 knockdown induced marked reduction in blastema size, neoblast cell numbers and proliferation

After observing that *tip60* knockdown affects blastema formation in planarians, our next step involved quantifying the blastema size in the amputated head, tail, and trunk regions, to examine the repressive impact of *tip60* knockdown on blastema formation. By evaluating the differences in blastema size, we aimed to understand the extent to which gene knockdown influences the regeneration process across different body regions. For this, we quantified the size of blastema formed in these amputated regions by calculating the ratio of the unpigmented area to the total area of the worm and compared the results with the control animals. The assessment of blastema size at both 5 DPA and 7 DPA revealed a notable decrease in blastemal growth (**Figure 4A**), indicative of impaired homeostasis and regeneration process, likely attributable to the knockdown of the *tip60* gene, and underscores its critical role in the normal regenerative processes of planarians.

**Fig. 4.**
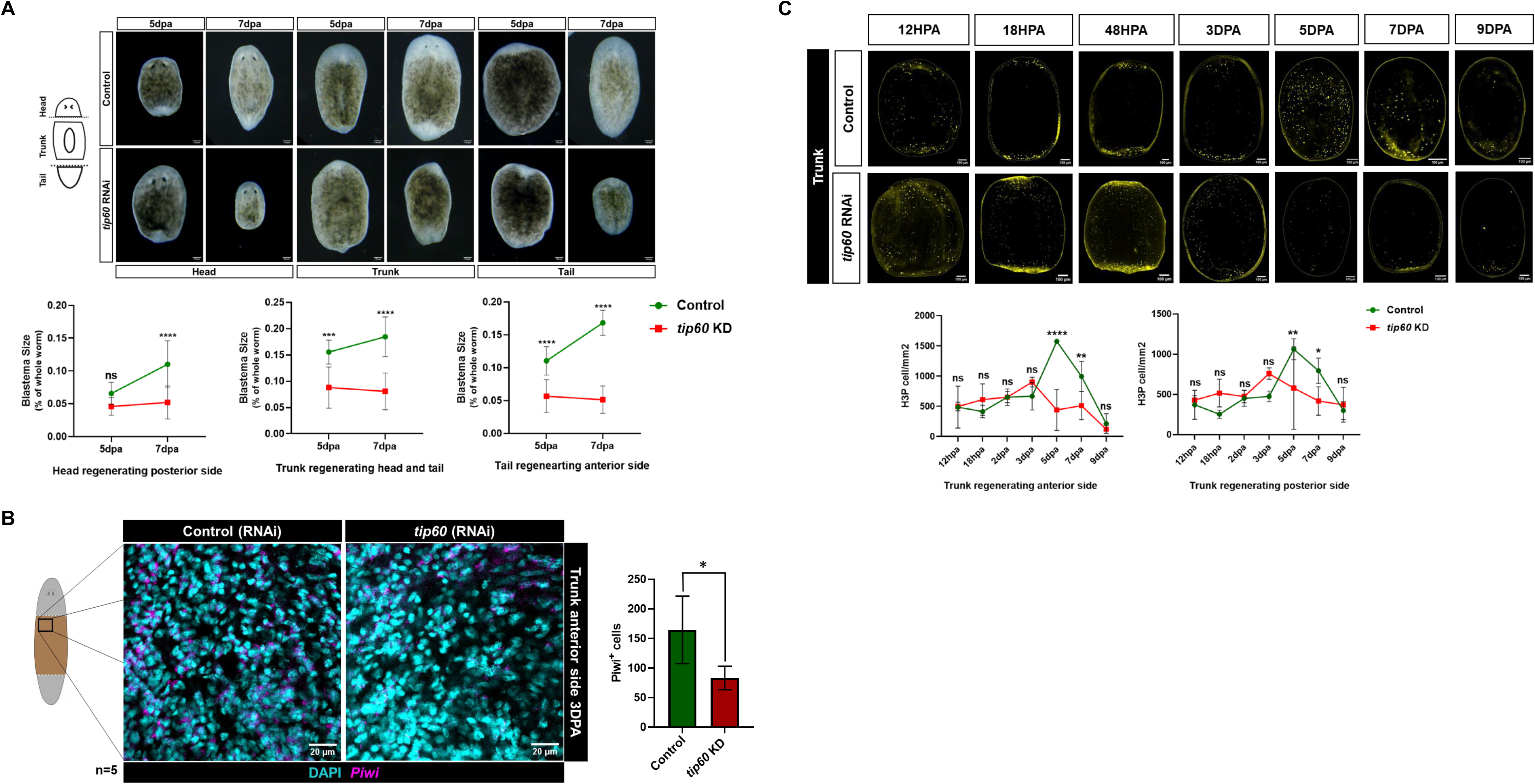
SMED-TIP60 plays a crucial role in regulating stem cell proliferation and maintenance. (A) Planarians were amputated transversely as shown in the cartoon image. Dark-field images of regenerating control and *Smed-tip60* RNAi planarians at 5 DPA and 7 DPA. Quantification of blastema size compared to whole planaria size was calculated for control and *Smed-tip60* RNAi planarians shown in the below panel for the head (posterior side), Trunk (anterior and posterior side), and tail (posterior side). (B) Single RNA fluorescence in situ hybridization (RNA-FISH) for *Smed-piwi-1* (neoblast cells maker) in trunk regenerating control and *Smed-tip60* RNAi planarians. Images shown here are single z-stack. (n=5, scalebar=20µm). Quantitation of piwi1+ cells in control and *tip60* RNAi condition is shown below. (C) Immunostaining of Histone 3 serine 10 phosphorylation (H3P) cells was performed for control and *Smed-tip60* RNAi planarians using anti-H3P antibody. Maximum intensity images showing the expression of H3P+ cells at different regeneration time points (12HPA, 18HPA, 48HPA, 3DPA, 5DPA, 7DPA, and 9DPA). (n=5, scale bar = 200µm). Quantification for H3P+ cells per mm^2^ is provided on the right side. All statistical analyses were performed using the graph pad Prism 8. Statistical test for Figures (A) and (C) is Sidak’s multiple comparison test, and Welch’s *t*-test for Figure (B). The error bar shows the standard deviation. Here, ns signifies p-value > 0.05, and *, **, ***, **** denotes p-value < 0.05, 0.01, 0.001, 0.0001.

Since the process of blastema formation in planaria is closely linked to the proliferation and differentiation of neoblast cells ^78^, we next wanted to evaluate the effect of TIP60 suppression on neoblast cell population. To visualize and quantify the number of neoblasts present in the planarian tissue, we performed RNA *in situ* hybridization for *Smedwi-1*, a well-established marker that specifically labels neoblasts. Our findings revealed a significant reduction in neoblast cell numbers in *Smed-tip60* KD animals compared to control animals at 3 DPA **(Figure 4B)**. Next, to determine if the reduction in neoblast numbers was due to a decrease in cell proliferation, we analyzed the proliferative status of neoblast cells by quantifying Histone 3 phosphorylated (H3P+) cells, which marks the neoblast cells specifically in the G2/M phase of the cell cycle. To track temporal changes in proliferative activity of neoblasts, immunostaining with H3P antibody was performed at different time points during the regeneration process, and presence of H3P+ cells were examined in *tip60* knockdown (KD) animals during head, trunk, and tail regeneration at various time points: 12 HPA (hours post-amputation), 18 HPA, 2 DPA, 3 DPA, 5 DPA, 7 DPA, and 9 DPA **(Figure 4C and Supp. Figure 4A)**. Notably, a significant reduction in H3P+ cells was observed in head-regenerating fragments, in *tip60* (RNAi) animals. In trunk-regenerating animals, the control group exhibited a mitotic peak at 5 DPA, while in *tip60* KD animals, this peak was sustained only until 3 DPA, followed by a marked decline in H3P+ cells, suggesting TIP60’s importance in sustaining neoblast proliferation beyond the initial stages. Similarly, in the anterior region of tail-regenerating animals, there was a notable decrease in H3P+ cells in *tip60* KD animals after 3 DPA. Together, these results suggest that TIP60 is vital for the sustained proliferation of neoblasts during different stages and regions of planarian regeneration highlighting the critical role of TIP60 in ensuring successful regenerative processes.

### SMED-TIP60 participates in regeneration program in planaria

After identifying the detrimental effects of *tip60* knockdown on normal physiological maintenance and regenerative processes, we wanted to examine the expression levels and localization pattern of the TIP60 protein during blastema formation in wild-type planaria post-amputation, aimed to further elucidate the role of SMED-TIP60 in planarian regeneration. For this we collected blastema samples from different regions (anterior and posterior side of the regenerating trunks) at various time points (12 HPA, 1 DPA, 2 DPA, 3 DPA, 5 DPA, 7 DPA, and 9 DPA), following amputation. Using Western blot analysis, we quantified the TIP60 protein levels to understand its role in blastema formation. Our results indicated that the expression level of the SMED-TIP60 protein started increasing immediately after amputation and reached maximum by 5 DPA, after which it started to decline **(Figure 5A)**. The increasing levels of the protein until 5 DPA indicate that it may be essential for initiating and supporting the early stages of blastema formation, such as cell proliferation, differentiation, and tissue remodelling. However, the subsequent decline in TIP60 expression after 5 DPA suggests that, once the initial phases of blastema formation are completed, the protein’s function may be downregulated which could be a regulatory mechanism designed to prevent excessive or uncontrolled growth.

**Fig. 5.**
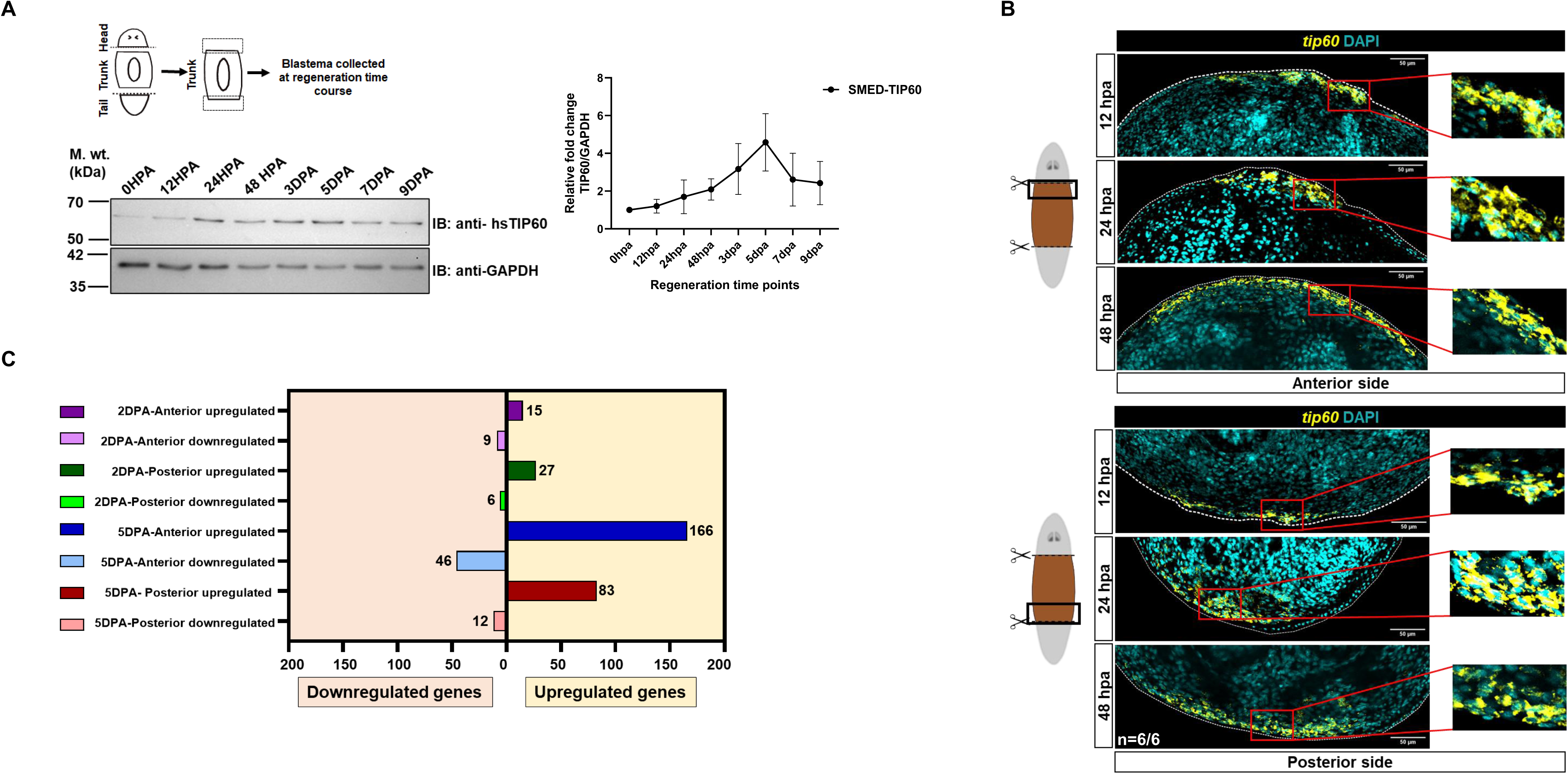
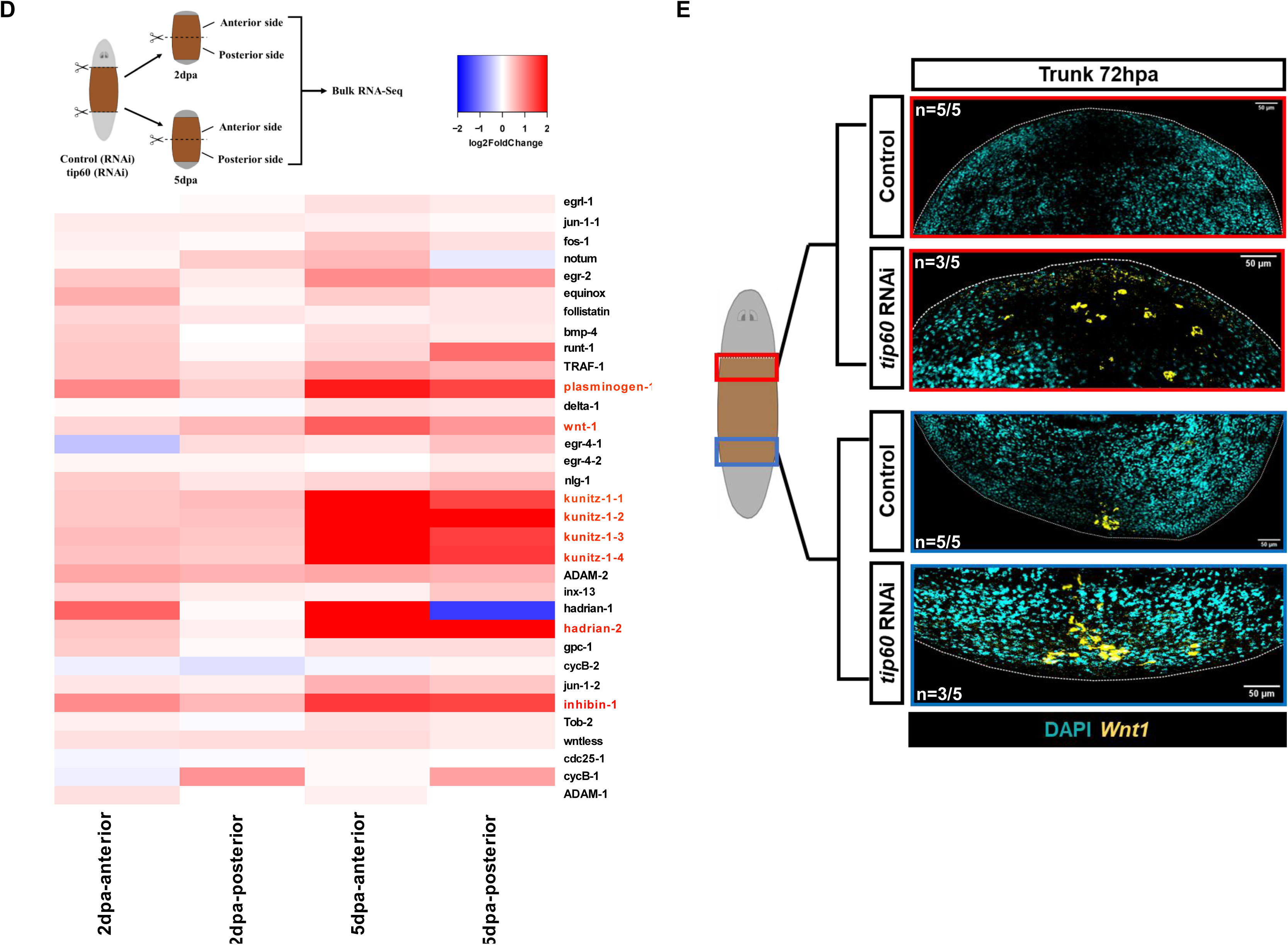
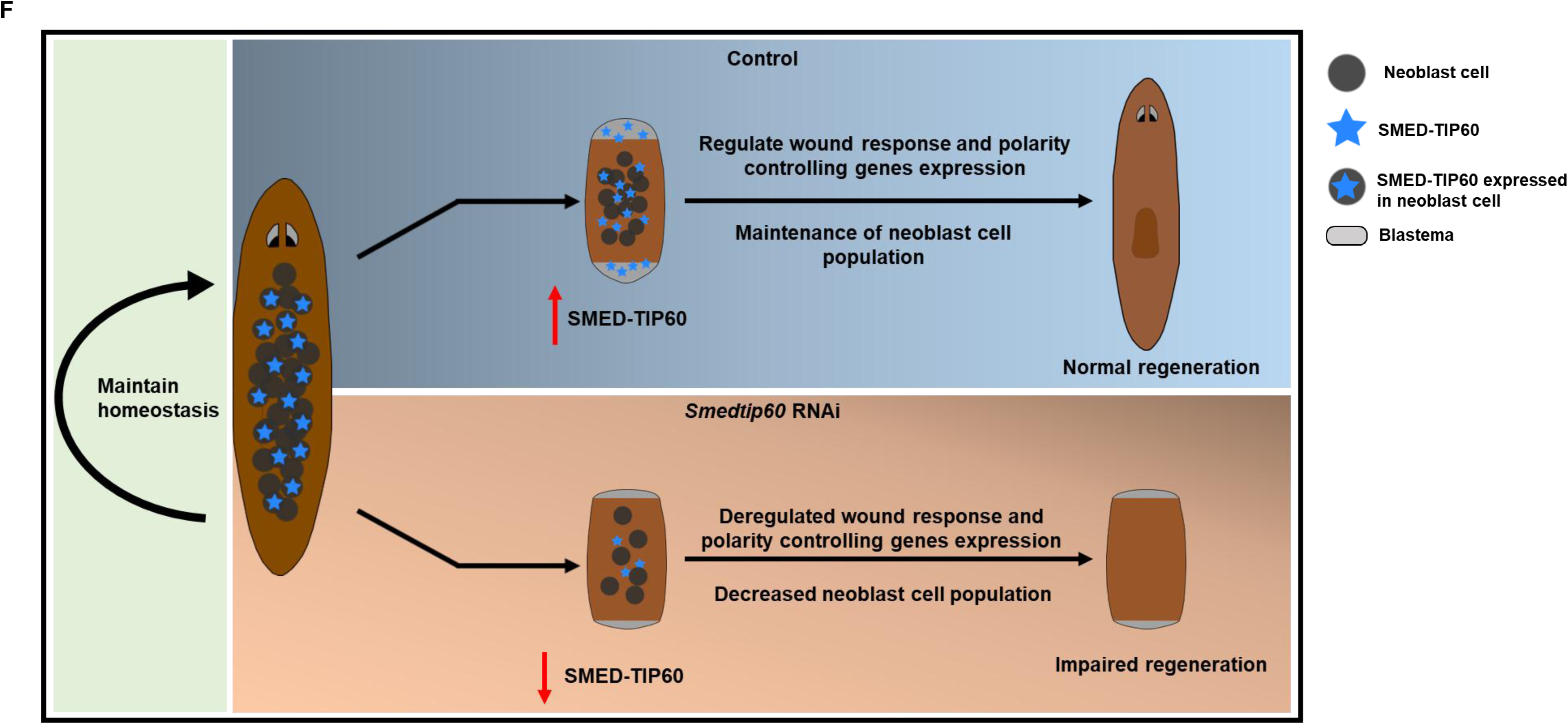
SMED-TIP60 regulates the regeneration process by controlling the wound response gene expression. (A) Wild-type planarians were amputated transversely and blastema of the trunk was collected as shown in the cartoon image for 12HPA, 24HPA, 48HPA, 3dpa, 5dpa, 7dpa, and 9dpa regeneration time points, and 0HPA was taken as a control. Collected samples were resolved on the SDS-PAGE and proceeded for Immunoblot using indicated antibodies. The intensity of the SMED-TIP60 bands was quantified using ImageJ and the graph was plotted after normalizing the intensity with GAPDH (internal control) shown on the right side. The graph was plotted for the average of three independent experimental replicates. The error bar shows the Standard error of the mean (SEM). (B) RNA in situ hybridization of *Smed-tip60* in wildtype planarians showing the expression of *Smed-tip60* at the anterior side (Top panel) and posterior side (Bottom panel) blastema during regeneration time points at 12HPA, 24HPA, and 48HPA. Maximum intensity images are shown here. (n= 6, scale bar = 50µm). (C) Graph showing the differential expression of genes at the 2 DPA and 5DPA (Anterior and posterior side). (D) Heatmap showing the expression for wound response genes at 2dpa and 5dpa for anterior and posterior side respectively from transcriptome profiles. Genes highlighted in red are the ones significantly expressed condition (E) Images showing the maximum intensity of Single RNA FISH for *wnt1* at 3 DPA anterior side (Red box) and 3 DPA posterior side (Blue box). n=5, scale bar=50µm. (F) Working model showing how TIP60 regulates the regeneration process in planaria by regulating the wound response genes expression.

Once we quantified and analyzed the expression level of TIP60 protein in blastema formation at the different time points after amputation, we wanted to determine if the TIP60 protein specifically expresses at the blastema and whether its expression varies within the blastema-specific site across different time points, after amputation. For this performed RNA *in situ* hybridization experiments to examine the expression of *tip60* in the anterior and posterior side of the regenerating trunks. Our analysis revealed that *tip60* is expressed at blastema site **(Figure 5B)** during regeneration suggesting its active involvement in the healing process. The expression of *tip60* was detected at the anterior and posterior stumps as early as 12 hours of regeneration and interestingly increased *Smed-tip60* increased accumulation localized near the outer edge or boundary of the newly forming tissue (blastema) during regeneration at both the anterior and posterior stumps **(Figure 5B)**. The early and increasing presence of *tip60* indicates its role in the initial wound response that could be important for providing positional information and coordinating tissue formation during regeneration.

To examine the gene expression changes because of *tip60* knockdown that contribute to regeneration defects in planaria, we conducted RNA sequencing (RNA-Seq) on regenerating *tip60* KD planarians. We collected samples at two critical time points: 2 days post-amputation, when TIP60 expression exhibited a two-fold increase, and 5 days post-amputation, when TIP60 expression peaked. For comprehensive analysis, we collected samples from both the anterior and posterior blastema ends of the regenerating trunk. Bulk RNA-Seq data analysis revealed **24** and **212** differentially expressed genes in *tip60* (RNAi) regenerating animals at the anterior end of the trunk at 2 DPA and 5 DPA, respectively, with a False Discovery Rate (FDR) ≤ 0.01 and a log fold change ≥ 1 **(Figure 5C**). While we identified **33** and **95** differentially expressed genes in *tip60* (RNAi) regenerating animals at the posterior blastema of the trunk at 2 DPA and 5 DPA, respectively, under the same FDR and fold change criteria **(Figure 5C)**. We conducted a comparative analysis of gene expression between 2- and 5-days post-amputation (DPA) within the anterior and posterior regenerating regions. Among the differentially expressed genes, 10 genes were commonly upregulated in the anterior region at both 2 DPA and 5 DPA, while only 1 gene was shared between the upregulated gene sets in the posterior region at these time points (**Supp. Figure 5A**). For downregulated genes, 3 were shared in the anterior region and 2 in the posterior region between 2 DPA and 5 DPA. We also compared gene expression between anterior and posterior regions at each time point. At 2 DPA, 4 genes were upregulated and 2 downregulated between the anterior and posterior ends, whereas at 5 DPA, 70 genes were upregulated and 4 downregulated (**Supp. Figure 5B**). These findings indicate increasing transcriptional divergence between anterior and posterior regions over time. Notably, in *tip60* (RNAi)-treated animals, the gene expression profiles at 2 and 5 DPA were distinct in the anterior and posterior regenerating tissues, suggesting limited functional overlap of SMED-TIP60 between these regions. Additionally, we examined the expression of known wound-induced genes across all four conditions (2 DPA-anterior, 2 DPA-posterior, 5 DPA-anterior, and 5 DPA-posterior) to further characterize region- and time-specific transcriptional responses during regeneration.

Since, SMED-TIP60 protein showed wound-induced increase indicating its involvement in initial stages of wound-response program, we also performed wound-induced genes expression analysis in *Smed-tip60* RNAi animals. Our analysis showed significant upregulation of some of the wound induced genes like *plasminogen-1, wnt-1, kunitz-1:4, hadrian-2 and Inhibin-1* in 5 DPA as compared to 2 DPA **(Figure 5D)** suggesting a failure in the down regulation of wound induced genes in the *tip-60* KD animals. Among the genes that demonstrated significant upregulation, the most notable is *wnt1*, for its critical role in determining polarity during the regeneration process. *wnt1* is an early wound-response gene that establishes body axis polarity during planarian regeneration ^79^. It is expressed by muscle cells at the dorsal midline of the posterior tip in intact planaria, acting as a posterior pole organizer. *wnt1* expression initiates approximately 6 hours post-amputation (HPA) and is subsequently suppressed by its antagonist *notum* between 24-30 HPA, a process crucial for head regeneration ^80^. To examine the temporal expression of *Wnt* we performed RNA in situ analysis for *wnt1* and the transcript levels were compared to the expression and localization patterns observed in control (RNAi) animals. In our analysis, we observed ectopic expression of *wnt1* in *tip60* (RNAi) animals **(Figure 5E)**. Notably, *wnt1* expression, which is typically absent at the anterior end in regenerating blastema, was detected at the anterior pole at 24 HPA compared to the control **(Supp. figure 5C)**. Furthermore, *wnt1*, which normally localizes specifically to the posterior pole, showed increased, widespread, and prolonged expression at the posterior end depicted by the increased immunofluorescence in the regenerating region of the blastema **(Figure 5E)**. This expression pattern was monitored until 3 DPA, revealing that KD animals displayed extended pattern of *wnt1* expression. The presence of *wnt1* at the anterior pole and its abnormally increased and widespread expression at the posterior side of the amputated trunk fragment in *tip60* knockdown (KD) animals suggest a prolonged wound response, thereby disrupting the balance of signalling pathways necessary for proper regeneration. This also suggests that TIP60 plays an important role in regulating the wound response by modulating the expression of *wnt1* critical for the anterior-posterior pole formation during regeneration. In conclusion, our findings show that *tip60* knockdown planarians failed to survive and exhibited abnormal and prolonged expression of wound-induced genes. This highlights the critical role of TIP60 in regulating the early wound response required for proper tissue regeneration in planarians **(Figure 5F)**.

## DISCUSSION

Planaria possess a remarkable regenerative capacity, enabling them to rebuild any lost tissues, organs, or body parts in response to injury or amputation. In case of injury, planaria immediately responds by closing the wound with a thin film of epidermis aided by muscle contraction. However, when the injury results in tissue loss, it triggers a wound response program, followed by a neoblast-mediated regenerative process to regrow the lost body parts ^36,81–84^. Any injury, irrespective of tissue loss, triggers a generic wound response by transcriptionally activating wound-induced genes within 0-24 hours post-injury ^84,85^.

The duration of activation of the first wave of wound-induced genes varies depending on the type of injury. In cases involving tissue loss, this initial wave is followed by a second wave of regenerative responses. During this phase, tissue patterning factors, particularly those associated with muscles, and genes linked to specialized neoblasts are activated ^85^. Achieving a balance between halting the general wound response program and activating the regeneration-specific program is necessary for the effective and successful restoration of missing tissues and the proper establishment of tissue patterning. However, the regulatory mechanisms behind these synchronized transcriptional changes during regeneration are not well understood. Various mechanisms, such as epigenetic modifications, chromatin remodelling, and posttranslational modifications, influence gene expression by modulating transcription. While research on the role of histone modifications and epigenetic modifiers in regulating gene expression in *planaria* remains limited, existing studies have underscored the importance of histone modification-mediated epigenetic regulation in these organisms ^86–88^. TIP60 is a multifunctional protein, which plays critical role in the maintenance of genomic integrity by participating in DNA damage repair process and mitosis, besides contributing to other important cellular process including transcriptional regulation, apoptosis, autophagy, and nuclear receptor signalling ^89^.

In mammals, TIP60 is identified as critical regulator for embryonic survival of mice as the ablation of the *tip60* gene (homozygous-null *TIP60*−/−) causes early embryonic lethality at the blastocyst stage. This is evidenced by the failure of cells in TIP60-null blastocysts to undergo hatching and sustain survival in culture ^90^. These findings underscore the essential role of TIP60, suggesting that its functions are non-redundant and cannot be compensated by other MYST family members. Further, it has been shown that TIP60 plays a pivotal role both in regulation of embryonic stem cells (ESCs) self-renewal during pre-implantation & differentiation during post-implantation ^91,92^. Similarly, the conditional knockout of *tip60* in murine hematopoietic stem cells (HSCs) exhibit critical role of TIP60 protein for both foetal and adult HSCs maintenance ^15^. Our findings in planaria showed that the depletion of *Smed-tip60* causes lethality in the intact animals as well as disrupts the formation of blastema, that consequently impairs the regeneration process (**Figure 3D and 3E**). Our findings are consistent with a recent study that demonstrated the impact of *tip60* knockdown on homeostasis and the regeneration process in planaria ^93^. However, it is noteworthy that the timing of the observed effects following knockdown differs significantly between the two studies. While the exact reason for this discrepancy remains unclear, it may be attributed to the differences in RNAi methodologies. In our hands, we were unable to achieve effective *tip60* knockdown using the feeding method and instead had to resort to dsRNA injection to obtain reliable results. The blastema formation is a hallmark of regeneration in planaria. It’s a cluster of undifferentiated cells that forms at the site of injury and is essential for regrowth of tissues and is considered central component of regenerative process that occur by epimorphosis ^94^. Additionally, we also found that the inability to form blastema in *tip60* RNAi animals correlated with a decrease in the total number of neoblast cells **(Figure 4B)**. Hence, the inability of *tip60* RNAi worms to form a blastema indicates that TIP60 is essential for the early stages of regeneration, and the depletion of TIP60, could disrupt these processes and affect signalling pathways, thereby preventing the self-renewal of neoblasts at the injury site leading to impaired regeneration.

The rise in TIP60 expression beginning around 24 hours post amputation (HPA) within the blastema and reaching its peak by approximately 5 DPA, indicates its sensitivity to injury. In planaria, the blastema is formed from proliferating neoblasts near the wound ^82,83^. TIP60 depletion not only reduces neoblast numbers but also halts the blastema formation at both the anterior and posterior ends **(Figure 4A and 4B)**. Our bulk RNA-seq data from regenerating head and tail fragments show alterations in transcriptional programs in the absence of TIP60, with these changes becoming more pronounced as TIP60 levels increase. These alterations are distinct at both the regenerating ends. The number of differentially expressed genes (DEGs) at 2 DPA in *tip60* knockdown is relatively low, but these DEGs are mostly unique to the regenerating head and tail ends. However, by 5 DPA, the number of DEGs increases nearly 8.8-fold at the head regenerating end, while at the tail regenerating end, there is only a 2.8-fold increase. This rise in DEGs at 5 DPA at both the ends correlates with the increased expression of SMED-TIP60 protein following injury **(Figure 5D)**. TIP60, by acetylating histones, can decompact the chromatin structure, and as a result is generally considered to act as a co-activator of gene transcription ^95,96^. However, contrary to this notion, our RNA-seq data revealed that most of the differentially expressed genes (DEGs) were upregulated at both the head (1.66-fold at day 2 and 3.6-fold at day 5) and tail (4.5-fold at day 2 and 6.9-fold at day 5) regenerating ends on both days of examination. Interestingly, similar observation was made in ESCs where in TIP60 knockdown condition out of 802 DEGs, 674 were upregulated while only 128 were down regulated ^91^. TIP60 has been identified as a chromatin modifier and remodeler in various model systems, where it has been shown to acetylate histones H2A, H3, and H4, and to facilitate the exchange of H2AX/v, leading to alterations in chromatin structure ^13,19,97^. While the overall sequence similarity between the planarian and human homologs of TIP60 was found to be limited, the planarian TIP60 retains key conserved domains and catalytic activity like that of as that of human TIP60. Higher sequence and structural similarity between catalytic MYST domain of planaria and human proteins, coupled with evidence from *in vitro* biochemical assays, confirms that SMED-TIP60 is a bona fide acetyltransferase protein capable of acetylating histones (**Figure 2C and 2D**). Thus, it would be interesting to investigate in future studies whether TIP60’s catalytic activity is essential for the observed alterations in gene expression, or if this function operates independently of its enzymatic activity.

Our *in-situ* hybridization analysis demonstrated that TIP60 is predominantly expressed at the periphery of both the anterior and posterior blastema as early as ∼12 hours post-amputation (HPA). This temporal and spatial expression pattern implies its potential role in wound response and blastema patterning. To substantiate this, we re-analyzed our RNA-seq dataset with a less stringent cutoff, uncovering dysregulated expression of several wound response-associated genes, expressed by both neoblasts and differentiated cells, encompassing W1, W2, and W3 categories corresponding to wound-responsive differentiated cells, and category W4 representing genes expressed in neoblasts ^83^. Several genes including *wnt1*, *notum*, *wntless*, which are involved in the decision-making process for head and tail regeneration, along with BMP-4 required for patterning of dorso-ventral axis, exhibited derestricted expression in the absence of SMED-TIP60 **(Figure 5D)**. *In situ* hybridization analysis of wnt1 in Smed-TIP60 RNAi animals revealed delocalized and abnormal expression at both the anterior and posterior poles **(Figure 5E)**. Additionally, Equinox, a gene expressed in wound epidermis immediately after injury and necessary for blastema formation at both ends ^73^ showed a biased expression at the anterior pole. Similarly, *runt-1* expressed in neoblasts near the wound and vital for promoting specification of neuron and eye precursor neoblasts ^83,98^ displayed enhanced expression at the posterior end at 5-day pi in SMED-TIP60 depleted condition. These findings indicate that Smed-TIP60 is required to maintain the wound-response genes program and to reset the positional information needed for regenerating lost or missing tissues.

Recently, during wound induced condition, human TIP60 is shown to regulate genes involved in actin reorganization and filopodia formation ^99^. However, HsTIP60 is able to execute these functions in cooperation with nuclear receptor PXR through its NR box at C-terminus. Interestingly, SMED-TIP60 lack NR box and it aligns with the absence of any nuclear receptor transcription family proteins detected in planaria to date. Further, in our comparative analysis, of SMED-TIP60 and its human homolog, we observed a noteworthy divergence in the homology of their chromodomains. While MYST domain exhibited higher levels of similarity, the chromodomains of these proteins shared significantly less homology. Chromodomains are known to recognize specific histone modifications, and differences in their structure can lead to differences in their interactions with histones and other proteins ^100^. The significant structural differences, detected between human and SMED chromodomain, indicate that SMED-TIP60 proteins may have different *in vivo* binding specificities, and may interact with different histone marks or protein partners. It seems plausible that the NRB appeared in human TIP60 as part of an evolutionary adaptation, possibly to meet the demands of more complex regulatory networks in human cells. TIP60 is known to acetylate both histone and non-histone protein substrates, including many crucial transcription factors. It will be important identify the physiologically relevant *in vivo* substrate for SMED-TIP60 in future. Besides, the human TIP60 protein functions within a multiprotein complex to execute its catalytic activity on nucleosomes, *in vivo* ^31,101^. Whether the planarian homolog functions within a similar complex is unknown and presents an interesting direction for future research.

In summary, our research identifies a role of essential chromatin modifier in *S. mediterranea*, suggesting that, despite significant sequence divergence, the conservation of key functional domains ensures the preservation of essential biochemical activities across species. Notably, *tip60* knockdown planarians failed to survive, and its absence impairs blastema outgrowth and disrupts the wound response program (**Figure 5F**). The prolonged wound response observed in *tip60* knockdown animals likely interferes with the regeneration process, leading to defects in planarian regeneration. These findings indicate that TIP60 is essential for both the survival and proper functioning of the organism.

## Acknowledgements

We thank SNIoE for all the necessary infrastructure and resources. AP thanks ICMR for Senior Research Fellowship. BJ acknowledge DST-WOA-A fellowship and AG acknowledge SERB-TARE fellowship.

## Declarations

Ethics approval and consent to participate: Not applicable

## Availability of data and materials

All data generated or analysed during this study are provided within the manuscript or supplementary information files.

## Authors’ contributions

AP- data collection, analysis, illustrations and tables, writing original draft and editing, prepared figures and tables, editing of final draft; AA- RNA-seq data analysis and related illustrations, editing the final draft; BJ- conceptualization and design of the study, data analysis, writing original draft, editing and preparation of final draft; DP- project supervision, resources, data analysis, editing original and final draft; AG- conceptualization and design of the study, data analysis, resources, writing original draft and editing, preparation of final draft. All authors show the final draft and agree with it.

