## Supplementary file for "TIP60, a key lysine acetyltransferase, acts as a wound-induced factor essential for efficient wound response in planaria"

**Supplementary Information**

**Supplementary figure 1:** (A) Multiple sequence alignment for TIP60 protein homologs of human, *Saccharomyces cerevisiae*, and *Schmidtea mediterranea* showing conserved chromodomain highlighted in yellow, MYST domain highlighted in cyan, and NR box in green only present in humans*.* (B) The three-dimensional structure of full-length human TIP60, ScESA1, and SMED-TIP60 Chromodomain is in yellow, the MYST domain is in magenta, and the NR box is in red. The below panel shows the superimposition of human TIP60 in yellow with SMED-TIP60 in blue with an RMSD of 16.298Å, ScEsa1 in red, and SMED-TIP60 in blue with an RMSD of 27.055Å. (C) Domain map of SMED-TIP60 showing the location of two primer sets designed for semi-qPCR analysis. (D) Dot plot showing the enrichment of *Smedtip60* in 0 HPA, 18 HPA-anterior, 18 HPA-anterior-unc22, 18 HPA-posterior. While *Smedwi-1* shows enrichment in 6 HPA-calcein. (HPA- hours post amputation)

**Supplementary figure 2:** (A) Cloning of *Smed-tip60* in pET28(a) vector. Agarose gel image showing the amplified product of *Smed-tip60.* Right-hand panel: Agarose gel image showing the positive clones of pET28(a)-Smed-tip60 plasmid as the fall out was observed at the size of the Smed-tip60 marked by an arrow. (B) Pairwise sequence alignment of human tip60 and Smed-tip60. Highlighted lysine residues are the known and putative sites of acetylation in human TIP60. SMED-TIP60 shows two conserved sites at the MYST domain (K266 and K324) highlighted in red color. (C) Agarose gel image showing the positive clones of Smed-tip60 in pRS314 yeast expression vector. Clones were confirmed as the fall out was at the size of Smed-tip60 and further proceeded for Sanger sequencing. (D) Complementation assay of Smed-tip60 in yeast system by transforming pRS31-*Smed-tip60* in LPY3500 yeast strain. Along with that pRS314 (-) control, and pRS314ScESA1(+) control transformed. Streaked plates were kept at a recessive temperature of 25°C and a restrictive temperature of 37°C.

**Supplementary figure 3:** (A) cDNA of control and *Smed-tip60* RNAi planarians was used for the qRTPCR showing knockdown of Smed-tip60. (biological replicates=3, technical replicates=3)

**Supplementary figure 4:** (A) Immunostaining using H3p antibody for Head and Tail at regeneration time points (12 HPA, 18 HPA, 48 HPA, 3 DPA, 5 DPA, 7 DPA, and 9 DPA). Maximum intensity images are shown here. n=5, Scale bar = 20µm. Quantification for H3P+ cells per mm^2^ is provided on the right side. Sidak’s multiple comparison test was used for the analysis. The error bar shows the standard deviation. Here, ns signifies p-value > 0.05, and *, **, ***, **** denotes p-value < 0.05, 0.01, 0.001, 0.0001.

**Supplementary figure 5:** (A) Venn diagram showing the overlapping and unique set of differentially expressed genes at 2DPA and 5DPA in anterior and posterior side. (B) ) Venn diagram showing the overlapping and unique set of differentially expressed genes in anterior and posterior side at 2 DPA and 5 DPA. (C) Single RNA-FISH for *wnt1* at 12 HPA and 24 HPA. Maximum intensity projection showed for control and *tip60* RNAi planarians. The red box denotes the anterior side of the trunk, and the blue box indicates the posterior side (Scale bar=50µm, n=5).

**Supplementary table 1:** List of Primers used in this study.

**Supplementary table 2 :** Accession IDs for all TIP60 protein sequences included in the phylogenetic analysis.

**Supplementary Figure 1**

**(A)**

```
ScESA1          MSHDGKEEPGIAKKINSVDDIIIKCQCWVQK-----NDEERLAEILSIN--TRKAPPKFY	53
HsTIP60         --------------MAEVGEIIEGCRLPVLRRNQDNEDEWPLAEILSVK--DISGRKLFY	44
SMED-TIP60      ----------------MFNKLCEGVKLPVKLKD---SDIYHFAEILKARENKRLGTKEYY	41
                                 ...:    :  *       .*   :****. .     .   :*

ScESA1          VHYVNYNKRLDEWITTDRINLDKEVLYPKLKATDEDNKKQKKKKAT-NTSETPQDS----	108
HsTIP60         VHYIDFNKRLDEWVTHERLDLKKIQFPKKEAKTPTKNGLPGSRPGS-PEREVPASAQASG	103
SMED-TIP60      VHFDEYNKRLDDWVSEDRMKMHEMVFPTKKIHKDSSTASLTALVSTNDCSNSPFGEANAQ	101
                **: ::*****:*:: :*:.:.:  :  *   .  ..       .:    : * .

ScESA1          --LQDGVDGFSR---------------------------E------NTD-----------	122
HsTIP60         KTLPIPVQITLRFNLPKEREAIPGGEPDQPLSSSSCLQPNHRSTKRKVEVVSPATPVPSE	163
SMED-TIP60      E--AENVEYKIRLNSSTSS---------------SALKNED-----QMS-----------	128
                      *:   *                           :      : .

ScESA1          --------------------VMDLDNLNVQGIKDENISHEDEIKK-LRTSGSM---TQNP	158
HsTIP60         TAPASVFPQNGAARRAVAAQPGRKRKSNCLGTDEDSQDSSDGIPSAPRMTGSLVSDRSHD	223
SMED-TIP60      --------------------LVTMDDRNCFSYLQEPV--ANVLPQTG--SMMVTSHQEHQ	164
                                         . *  .  ::     : : .    :  :    .:

ScESA1          HEVARVRNLNRIIMGKYEIEPWYFSPYPIELTDEDFIYIDDFTLQYFGSKKQYERYRKKC	218
HsTIP60         DIVTRMKNIECIELGRHRLKPWYFSPYPQELTTLPVLYLCEFCLKYGRSLKCLQRHLTKC	283
SMED-TIP60      DIVTRMRNIDMIVLGKSRIKPWYFSPYPEELTELDAIFICEFCLKYFKSCFCLQRHLEKC	224
                . *:*::*:: * :*: .::******** ***    ::: :* *:*  *    :*:  **

ScESA1          TLRHPPGNEIYRDDYVSFFEIDGRKQRTWCRNLCLLSKLFLDHKTLYYDVDPFLFYCMTR	278
HsTIP60         DLRHPPGNEIYRKGTISFFEIDGRKNKSYSQNLCLLAKCFLDHKTLYYDTDPFLFYVMTE	343
SMED-TIP60      RLRHPPGNEIYRKNPLSFFEIDGRKNKKYAQNLCLLAKLFLDHKTLYYDTDPFLFYILCE	284
                 ***********.. :*********::.:.:*****:* **********.****** : .

ScESA1          RDELGHHLVGYFSKEKESADGYNVACILTLPQYQRMGYGKLLIEFSYELSKKENKVGSPE	338
HsTIP60         YDCKGFHIVGYFSKEKESTEDYNVACILTLPPYQRRGYGKLLIEFSYELSKVEGKTGTPE	403
SMED-TIP60      YDEEGFHIVGYFSKEKESNDDNNVACILTLPPYQRKGYGKLLIEFSYELSKLENKTGSPE	344
                 *  *.*:********** :. ********* *** *************** *.*.*:**

ScESA1          KPLSDLGLLSYRAYWSDTLITLLVEHQKEI------------------------------	368
HsTIP60         KPLSDLGLLSYRSYWSQTILEILMGLKSESGERP----------QIT-------------	440
SMED-TIP60      KPLSDLGLLSYRSYWCQTIMEVIIDVANSADGAANSSSSAFDPPSISLKYEFTSWKPLAT	404
                ************:**.:*:: :::   ..

ScESA1          --------TIDEISSMTSMTTTDILHTAKTLNILRYYKGQHIIFLNEDILDRYNRLKAKK	420
HsTIP60         ---------INEISEITSIKKEDVISTLQYLNLINYYKGQYILTLSEDIVDGHERAMLKR	491
SMED-TIP60      VFIYIYVCVCSDVVEKTSINPKDLNATLNTMNIIYYSKGQHVIVVTKKMLDKFKASMAKR	464
                          .:: . **:.  *:  * : :*:: * ***::: :.:.::* .:    *:

ScESA1          RRTIDPNRLIWKPPVFTASQLRFAW	445
HsTIP60         LLRIDSKCLHFTPKDWSKRG---KW	513
SMED-TIP60      KYRIDAKLLNWHPKDWSKRG---RW	486
                   ** : * : *  ::       *
```

Chromodomain, MYST domain, NR box

**(B)**

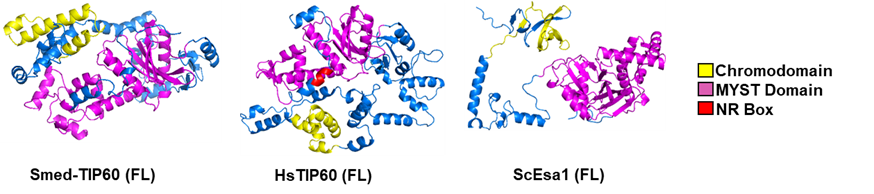

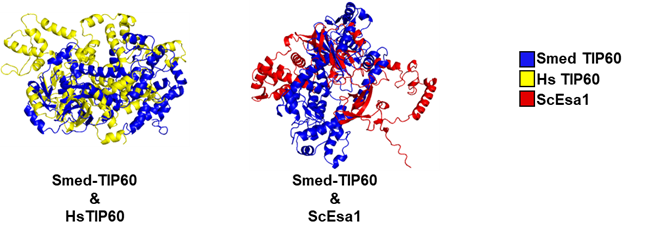

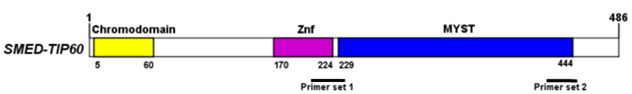
**(C)**

**(D)**

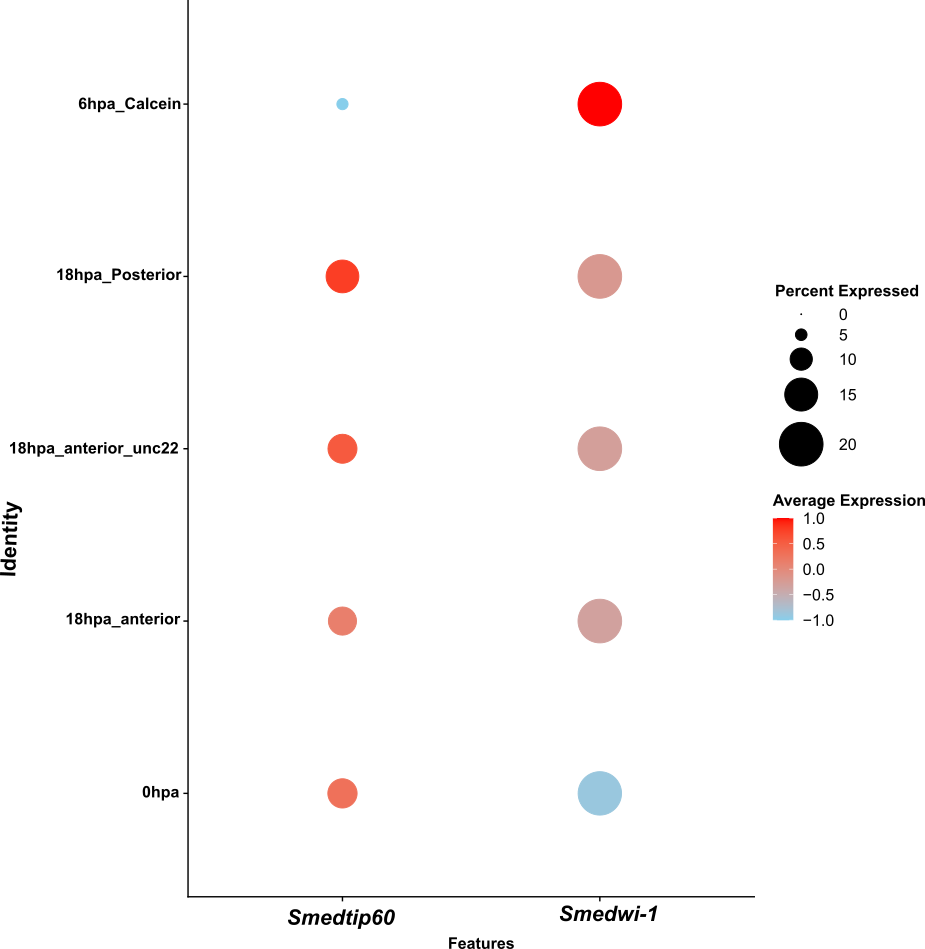
**Supplementary Figure 2**

(A)
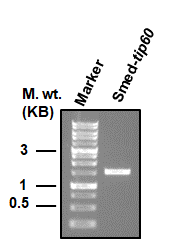

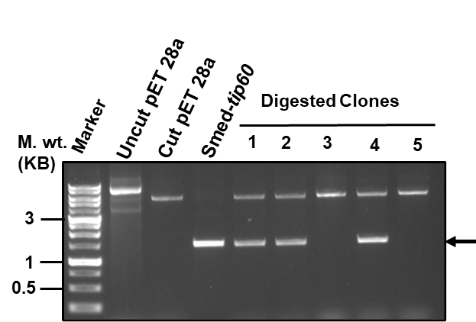

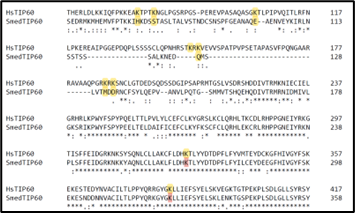
**(B)**

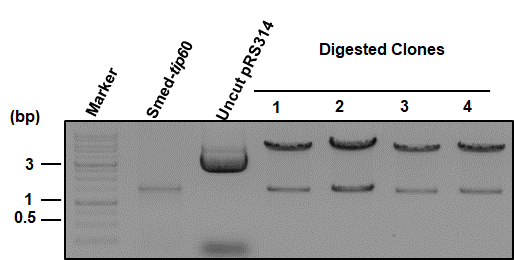
**(C)**

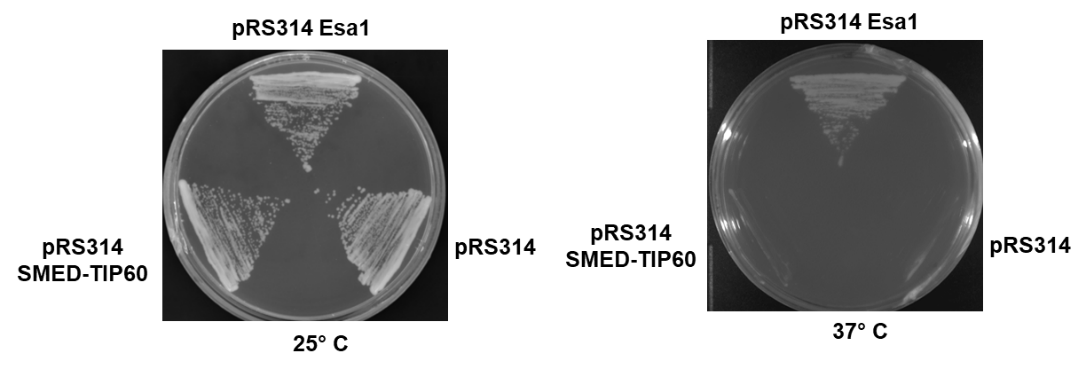
**(D)**

**Supplementary Figure 3**

**(A)**

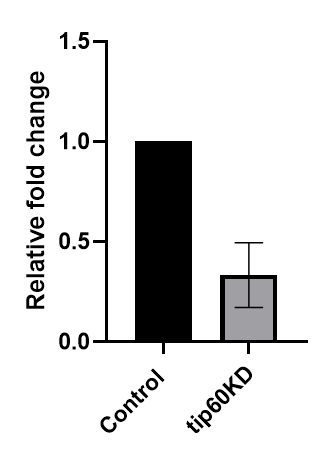

**Supplementary Figure 4**

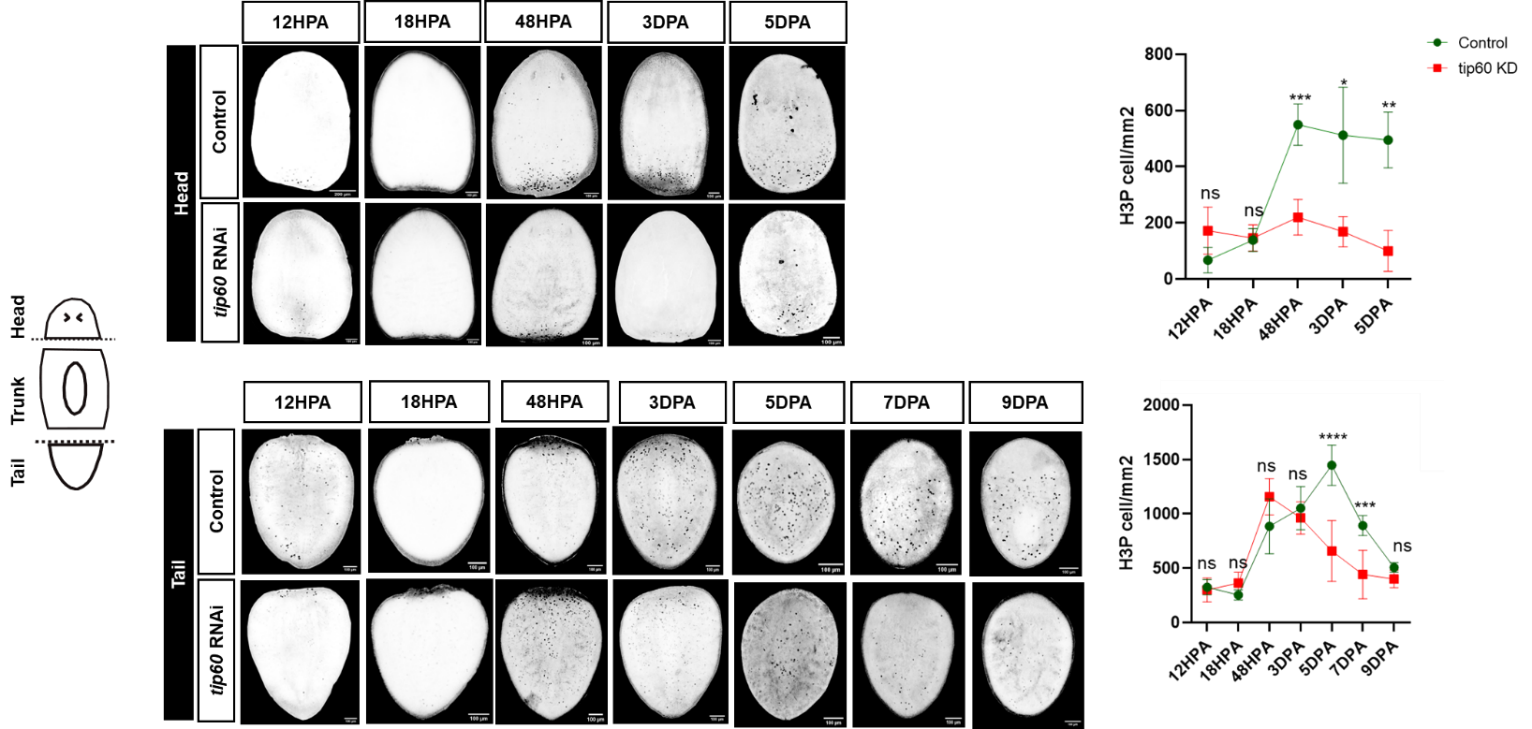
**(A)**

**Supplementary Figure 5**

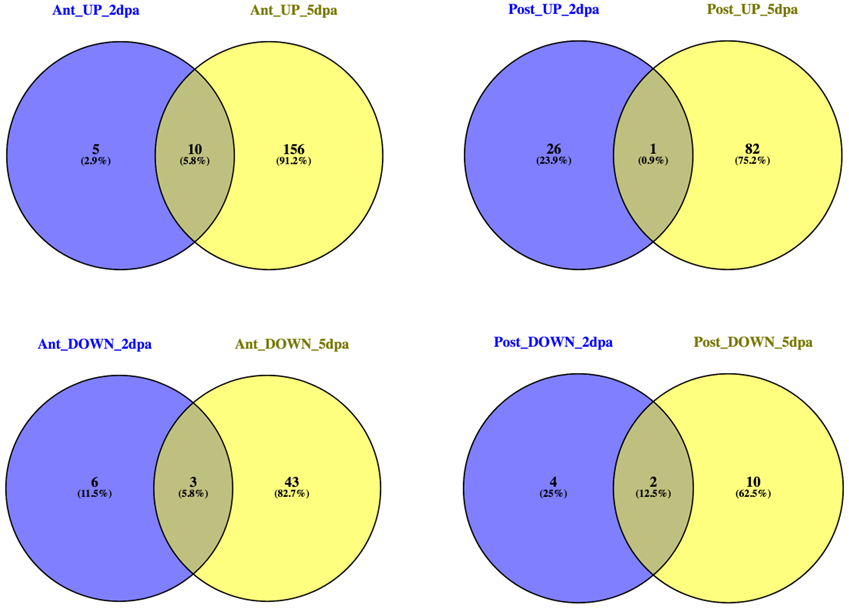
**(A)**

**(B)**
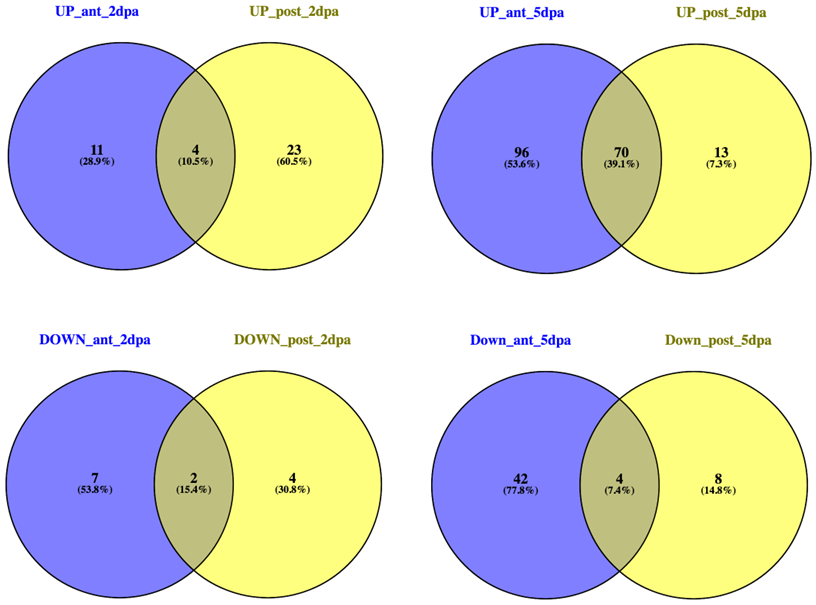

**(C)**

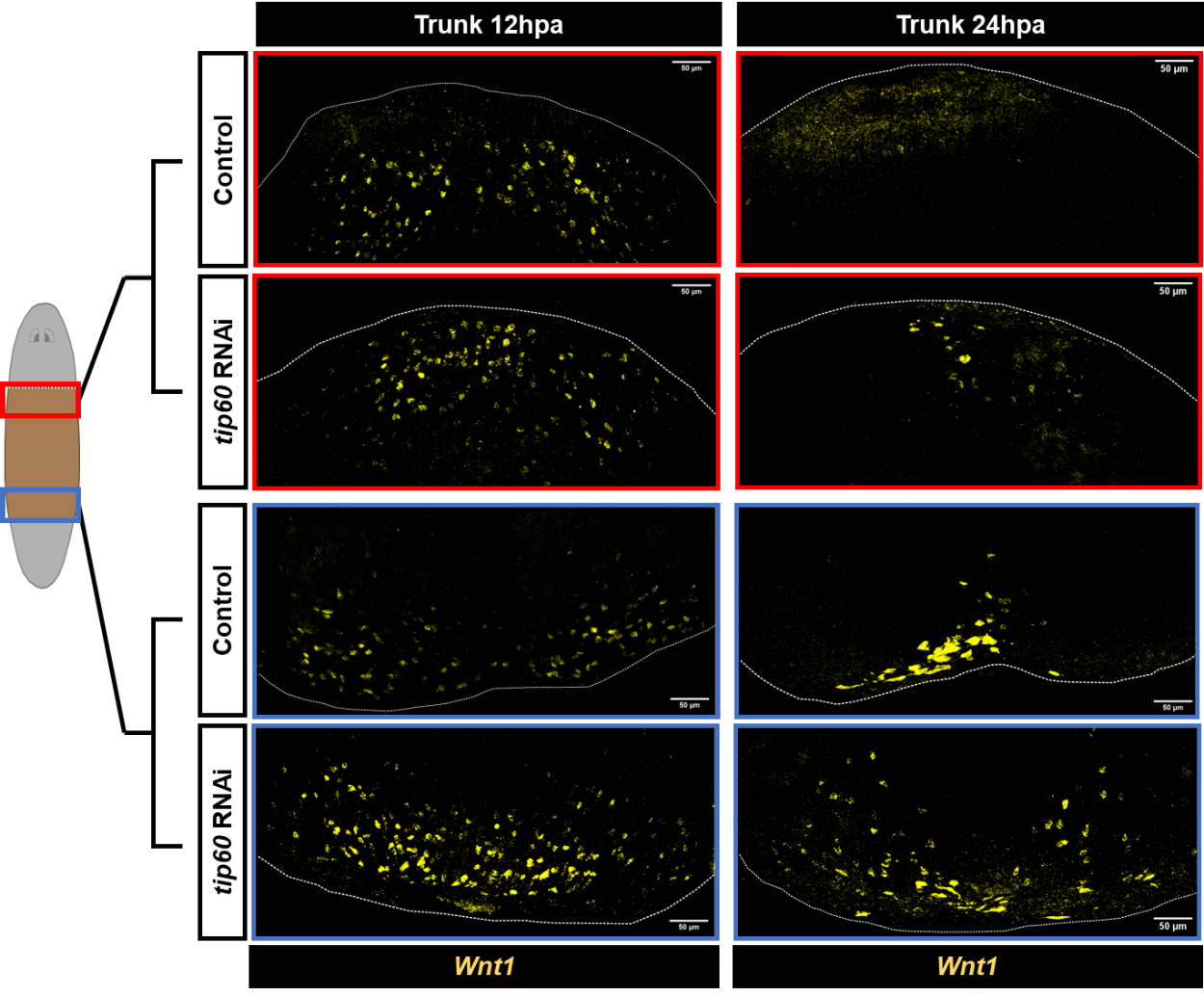

**Supplementary Table 1 :** List of primers used in the study

| **Used for study** | **Primer name** | **Primer sequence** |
| --- | --- | --- |
| *In vitro* HAT and Autoacetylation assay | SmedTIP60 FL fwd | CGGGATCCATGTTTAATAAGTTATGTG |
|  | SmedTIP60 FL rev | GCGTCGACTTACCAGCGACCTCGTTTGGACC |
| qRTPCR | SmedTIP60 fwd1 | TGCCCAAGAAGCGGAAAATG |
|  | SmedTIP60 rev1 | TATTGCGCATCCTAGTGACG |
|  | SmedTIP60 fwd2 | TTGGAAACCATTGGCAACGG |
|  | SmedTIP60 rev2 | ACCAATCCTTTGGATGCCAG |
|  | SmedActin fwd1 | CCCATTGAACACGGCATTG |
|  | SmedActin_rev1 | TGCATACAATGACAACACTGC |
|  | SmedActin fwd2 | GGAGGCTCAACAATGTATCC |
|  | SmedActin rev2 | TGGACCAGATTCGTCATATTC |
|  | SmedGAPDH fwd1 | CTGTTGAAGCCAAGGATGAC |
|  | SmedGAPDH rev1 | GGCAGAAGGAGCTGAAATG |
|  | SmedGAPDH fwd2 | AAAAGCTTCTGAAACTTCAATG |
|  | SmedGAPDH rev2 | GTAGGCAATCAAATCAACAAC |
| dsRNA/Probe | SmedTIP60 fwd | GCCCAAGAAGCGGAAAATGT |
|  | SmedTIP60 rev | ttgccgtaacctttccgttg |
|  | SmedTIP60 T7fwd | TAATACGACTCACTATAGGCCCAAGAAGCGGAAAATGT |
|  | SmedTIP60 T7rev | TAATACGACTCACTATAGTTGCCGTAACCTTTCCGTTG |
|  | SmedWnt1 fwd | TTCCATCTTTTATTCTCAGAGTTTG |
|  | SmedWnt1 rev | TTGATTGGATAAAAATGAGGAGTT |
|  | SmedWnt1 T7fwd | TAATACGACTCACTATAGTTCCATCTTTTATTCTCAGAGTTTG |
|  | SmedWnt1 T7rev | TAATACGACTCACTATAGTTGATTGGATAAAAATGAGGAGTT |
|  | Smedwi1 fwd | GTCTCAGAAAACAACTAAAGGTACAG |
|  | Smedwi1 rev | TGCTGCAATACACTCGGAGACA |
|  | Smedwi1 T7fwd | TAATACGACTCACTATAGGTCTCAGAAAACAACTAAAGGTACAG |
|  | Smedwi1 T7rev | TAATACGACTCACTATAGTGCTGCAATACACTCGGAGACA |

**Supplementary Table 2 :** Accession IDs for all TIP60 protein sequences included in the phylogenetic analysis.

**Protists -** *Acanthamoeba castellanii* - [XP_004344594.1](https://www.ncbi.nlm.nih.gov/protein/XP_004344594.1?report=genbank&log$=prottop&blast_rank=1&RID=5JF3MEGC013)

*Monoraphidium neglectum* - [XP_013906192.1](https://www.ncbi.nlm.nih.gov/protein/XP_013906192.1?report=genbank&log$=prottop&blast_rank=1&RID=5JF87NZ9013)

*Porphyridium purpureum* - [KAA8498568.1](https://www.ncbi.nlm.nih.gov/protein/KAA8498568.1?report=genbank&log$=prottop&blast_rank=1&RID=5JF91HB2013)

**Fungi -** *Neocallimastix californiae* - [ORY44253.1](https://www.ncbi.nlm.nih.gov/protein/ORY44253.1?report=genbank&log$=prottop&blast_rank=1&RID=5JFMKG98013)

*Glomus cerebriforme* - [RIA96599.1](https://www.ncbi.nlm.nih.gov/protein/RIA96599.1?report=genbank&log$=prottop&blast_rank=1&RID=5JFN3XK1016)

*Cryptococcus neoformans* - [OWZ57330.1](https://www.ncbi.nlm.nih.gov/protein/OWZ57330.1?report=genbank&log$=prottop&blast_rank=1&RID=5JFNMHK7016)

*Neurospora crassa* - [KAK3491619.1](https://www.ncbi.nlm.nih.gov/protein/KAK3491619.1?report=genbank&log$=prottop&blast_rank=1&RID=5JFV42GK013)

*Saccharomyces cerevisiae* - [AJT72155.1](https://www.ncbi.nlm.nih.gov/protein/AJT72155.1?report=genbank&log$=prottop&blast_rank=2&RID=5JFWGHA1013)

**Plantae** - *Physcomitrium patens* - [XP_024399982.1](https://www.ncbi.nlm.nih.gov/protein/XP_024399982.1?report=genbank&log$=prottop&blast_rank=2&RID=5JGA73UR013)

*Zea mays* - [AAM89290.1](https://www.ncbi.nlm.nih.gov/protein/AAM89290.1?report=genbank&log$=prottop&blast_rank=3&RID=5JG9WTVN016)

*Arabidopsis thaliana* - [NP_196536.1](https://www.ncbi.nlm.nih.gov/protein/NP_196536.1?report=genbank&log$=prottop&blast_rank=2&RID=5JG939ZC013)

**Animalia -** *Caenorhabditis elegans* - [NP_504796.1](https://www.ncbi.nlm.nih.gov/protein/NP_504796.1?report=genbank&log$=prottop&blast_rank=1&RID=5JG4CZGG016)

*Dugesia japonica* - [QDR71492.1](https://www.ncbi.nlm.nih.gov/protein/QDR71492.1?report=genbank&log$=prottop&blast_rank=1&RID=5JG3YWXX013)

*Drosophila melanogaster* - [NP_001259234.1](https://www.ncbi.nlm.nih.gov/protein/NP_001259234.1?report=genbank&log$=prottop&blast_rank=1&RID=5JG3B14A016)

*Nematostella vectensis* - [XP_032230595.1](https://www.ncbi.nlm.nih.gov/protein/XP_032230595.1?report=genbank&log$=prottop&blast_rank=2&RID=5JG2TW1X016)

*Danio rerio* - [XP_073806416.1](https://www.ncbi.nlm.nih.gov/protein/XP_073806416.1?report=genbank&log$=prottop&blast_rank=3&RID=5JG1MBDX016)
